# Enforced ZFP281 expression delays breast cancer initiation and can provide lifelong protection against breast cancer metastasis

**DOI:** 10.64898/2026.04.13.718280

**Authors:** Deepak K. Singh, Hongwei Zhou, Nyima Sherpa, Xiang Yu Zheng, Artem Lomakin, Pedram Razghandi, Xin Hunag, Rama Kadamb, Suryansh Shukla, Luis E. Valencia Salazar, David Entenberg, Deyou Zheng, Christina Curtis, Jianlong Wang, Julio A. Aguirre-Ghiso

**Affiliations:** Department of Cell Biology, Albert Einstein College of Medicine, Bronx, NY 10461, USA; Department of Oncology, Albert Einstein College of Medicine, Bronx, NY 10461, USA; Cancer Dormancy Institute, Albert Einstein College of Medicine, Bronx, NY 10461, USA; Gruss-Lipper Biophotonics Center, Albert Einstein College of Medicine, Bronx, NY 10461, USA; Institute for Aging Research; Albert Einstein College of Medicine, Bronx, NY 10461, USA; Montefiore Einstein Comprehensive Cancer Center, Albert Einstein College of Medicine, Bronx, NY 10461, USA; Ruth L. and David S. Gottesman Institute for Stem Cell Research and Regenerative Medicine, Albert Einstein College of Medicine, Bronx, NY 10461, USA; Marilyn and Stanley M. Katz Institute for Immunotherapy for Cancer and Inflammatory Disorders, Albert Einstein College of Medicine, Bronx, NY 10461, USA; Department of Medicine, Columbia Center for Human Development and Stem Cell Therapies, Herbert Irving Comprehensive Cancer Center, Columbia University, New York, NY, USA; Department of Genetics, Albert Einstein College of Medicine, Bronx, NY; Department of Medicine-Med/Oncology, Stanford University, Stanford, CA, USA; Department of Pathology, Cancer Dormancy Institute, Montefiore Einstein Comprehensive Cancer Center, Albert Einstein College of Medicine, New York; The Saul R. Korey Department of Neurobiology, Dominick P. Purpura Department of Neuroscience, Data Science Institute, Albert Einstein College of Medicine, Bronx, NY

**Keywords:** Breast cancer, Dormancy, ZFP281, Metastasis, DCIS, Mammary gland

## Abstract

Breast cancer metastatic reactivation and its links to mammary development are largely unknown. Here, using conditional knockout and overexpression in normal and HER2+ mammary glands, we show that the dormancy regulator ZFP281 promotes branching and dissemination while suppressing growth, and its overexpression can even override HER2-driven cancer initiation. Notably, while ZFP281 does not limit HER2-driven early dissemination, it constrains DCC plasticity, confining cells to a dormant mesenchymal/hybrid-like state and effectively suppressing metastasis throughout the mouse lifespan. ZFP281 is induced by estrogen, progesterone, and glucocorticoid signaling, and RNA sequencing of early lesions revealed that it regulates glutathione metabolism and ferroptosis, potentially supporting fitness during dormancy, while repressing angiogenesis, Th17-like inflammation, innate immune genes, and pro-inflammatory programs that might otherwise trigger awakening. Integrating these findings with human data, we show that DCIS and IBC (invasive breast cancer) lesions that later relapse are selectively enriched for ZFP281-regulated M-like and dormancy signatures and, after pregnancy, depleted for a ZFP281-induced stress-autophagy module, indicating that erosion or imbalance of these programs marks lesions that seed DCCs with higher reactivation potential. We propose that ZFP281 acts as a hormone-regulated dormancy gatekeeper that uncouples dissemination from growth, enforcing a metabolically fit, angiogenesis-low, immune-evasive dormant state in breast DCCs, thereby shaping the timing of metastatic relapse and potentially exploitable for durable prevention of metastasis.

**STATEMENT OF SIGNIFICANCE:** ZFP281, a hormone-regulated dormancy gatekeeper, uncouples dissemination from growth and constrains DCCs into long-term arrest, defining human gene signatures that distinguish dormancy-prone from awakening-prone lesions and predict breast cancer relapse dynamics.

## INTRODUCTION

Cancer patients predominantly succumb to the consequences of metastatic relapse, which originates from cancer cells that disseminate and can remain dormant for years or even decades before reactivating.^1^ However, our understanding of the intrinsic and microenvironmental programs that regulate dormancy and reactivation of disseminated cancer cells (DCCs) remains limited. We showed, using mammosphere studies and rigorous mouse models, that the Zinc finger protein 281 (ZFP281) drives a mesenchymal-like (M-like) dormancy program that controls the early dissemination of breast cancer cells and locks them in a dormant state by maintaining an M-like program in DCCs. ZFP281 inhibited the conversion of dormant DCCs into actively growing metastatic lesions by preventing the acquisition of an epithelial-like (Ep-like) proliferation program.^1^ We also showed that ZFP281 is induced by TGFβ2-TGFβRIII signaling activated by lung alveolar macrophages (AMs), and that loss of these AMs led to loss of ZFP281, dormant DCC awakening, and metastatic breakthrough.^2^

ZFP281 is a transcription factor that regulates stemness, pluripotency, and epigenome reprogramming during embryonic development by coordinating DNMT3 and TET1.^3–5^ In mouse embryonic stem cells, it blocks the transition from the primed to the naïve pluripotent state.^4^ ZFP281 is expressed early in embryogenesis in the undifferentiated trophoblast stem cell population in an FGF-dependent manner.^6^ It interacts with MLL or COMPASS subunits to occupy the promoter of its target genes and regulate their expression.^6^ ZFP281 is a crucial partner of OCT4 in embryonic stem cells (ESCs) and during the naïve-to-primed pluripotent state transition, where it reorganizes enhancer landscapes of lineage-specific genes such as *T, Otx2, Eomes and Gsc*.^5,7^ It also maintains and coordinates a balance between the functions of TET1 and TET2 in epigenomic configuration.^3^ ZFP281 physically interacts with Oct4, Sox2, and Nanog^8^ and multiple epigenetic pathways,^9,10^ regulating stem cell pluripotency and early embryogenesis. Mammary gland (MG) development uses similar gene programs and pathways as those activated during embryonic development.^11,12^ For example, pluripotency-associated signaling pathways and epigenetic regulators such as LIF/STAT3, WNT, FGF/FGFR, and TET2, which are necessary for embryonic development^3,4,6,13–16^, also play essential roles in MG development and physiology.^17–20^

These findings, coupled with the fact that ZFP281 is primarily an embryonic transcription factor (TF), were intriguing for several reasons: (i) ZFP281 expression is very low in resting wild-type MGs, (ii) it is upregulated by HER2 and PyMT early lesion (EL) phase (e.g., ductal carcinoma *in situ* (DCIS)) in mouse models but also irrespective of subtype in human DCIS samples, and (iii) it is silenced as primary tumors (human and mouse) and metastasis become highly proliferative. These observations and the fact that our prior data suggested a potential deregulation of mammary branching programs during early dissemination and dormancy^1,21^ led us to hypothesize that early breast cancer lesions may tap into ZFP281 to deregulate a branching morphogenesis program, leading to early dissemination of a persisting dormant DCC population that still carries inherent plasticity for reactivation and metastasis initiation. Here, we used conditional knockout and overexpression models in normal and HER2-positive mammary glands to reveal that ZFP281 indeed regulates hormone receptor-mediated branching morphogenesis. It facilitates branching programs while simultaneously silencing growth programs. The latter function is so powerful that when overexpressed, ZFP281 can override HER2-driven cancer initiation. Interestingly, ZFP281 does not limit early dissemination but constrains disseminated cancer cells to a dormant state, thereby suppressing metastasis throughout the mouse’s lifespan. Analysis of human gene signatures linked to relapse rates revealed that primary lesions associated with late relapse (10, 15, or 20 years) are indeed enriched for ZFP281-regulated programs, supporting the idea that these programs may control relapse dynamics in humans. DCIS lesions prone to relapse, sometimes more than a decade later, were also enriched in M-like, ZFP281-regulated signatures.

## RESULTS

### Over-expression of ZFP281 suppresses HER2-driven primary tumor development

To investigate the role of ZFP281 in tumorigenesis and metastatic progression, we developed mouse models with conditional knockout (cKO) and conditional overexpression (cOE) of ZFP281 in the luminal cells of MG **(S. Fig. 1A)**. Briefly, the MG-specific *Zfp281* cKO animals were generated by breeding C57BL/6 mice containing *Zfp281^f/f^* ^22^ with FVB mice expressing Cre recombinase under the control of the mouse mammary tumor virus promoter (MMTV-Cre).^23,24^ To develop the MG-specific *Zfp821* cOE mouse model, we first created a *Zfp281* cOE mouse model by inserting an extra copy of the *Zfp281* gene, whose expression is driven by the CAG promoter upon the removal of a floxed STOP cassette, into the H11 locus of C57BL/6NTac mice. We then bred C57BL/6NTac mice carrying *Zfp281^f/+^* with FVB MMTV-Cre to excise the STOP cassette and achieve the over-expression of *Zfp281* in mammary luminal cells. ZFP281 wild-type (WT: *Zfp281^+/+^*; MMTV-Cre), conditional knockout (cKO: *Zfp281^f/f^*; MMTV-Cre), and conditional over-expression (cOE: *Zfp281^f/+^*; MMTV-Cre; here f/+ represents 3 alleles of *Zfp281*) mice were verified by PCR genotyping and western blot analysis **(Fig. 1A-B).** Immunofluorescence (IF) for ZFP281 was performed in the MG tissue sections to confirm the conditional KO and OE of ZFP281 in luminal cells **(S. Fig. 1B)**. Luminal cells and basal and myoepithelial cells in the MG were specifically marked with Cytokeratin 8/18 (CK8/18) and Cytokeratin 5 (CK5) or Cytokeratin 14 (CK14), respectively.

**Figure 1.**
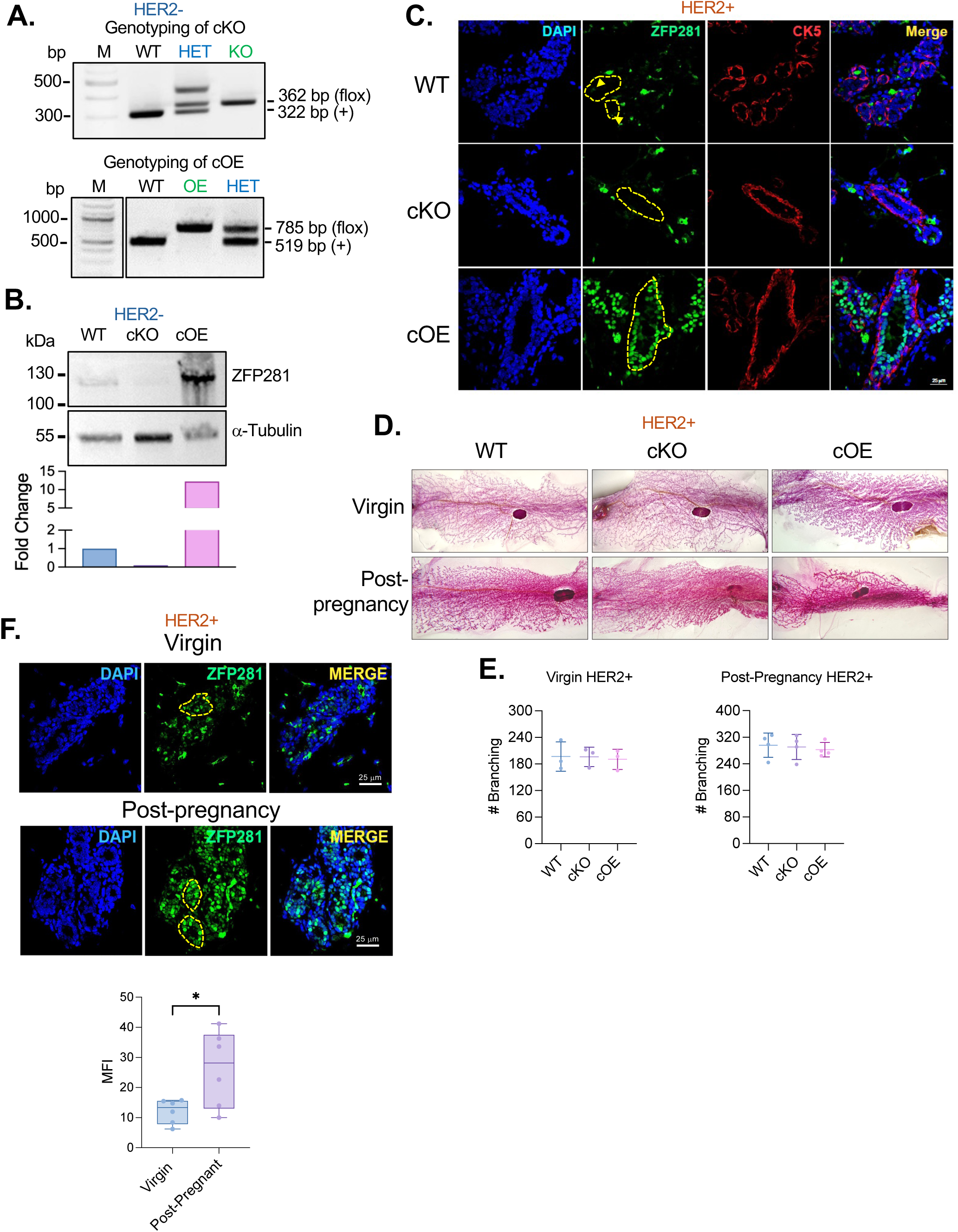
ZFP281 is induced during pregnancy. **A.** Confirmation of Zfp281 knockout and over-expression in the MMTV-Cre mice by PCR. **B.** Expression of ZFP281 in the organoids prepared from the mammary glands of 10-week-old mice. The graph shows densitometric quantification of ZFP281 expression. **C.** Immunofluorescence staining of ZFP281 in the mammary glands of MMTV-HER2 mice showing luminal cells specific knockout and overexpression of ZFP281. Basal cells in the ducts are stained for cytokeratin 5 (CK5). **D.** Whole mounts of inguinal mammary glands from virgin (10 weeks age) and post-pregnant (16-18 weeks age) mice showing no difference in the branching morphogenesis in ZFP281 KO and OE animals. **E.** Graphs show the quantification of branching. **F.** ZFP281 IF staining in the mammary gland of virgin and post-pregnant wildtype mice, showing induction of ZFP281 expression upon pregnancy. The graph shows the quantification of ZFP281 expression. MFI: Mean fluorescent intensity. Statistical test: Simple t-test, n=6.

Next, we bred *Zfp281^f/f^*; MMTV-Cre (cKO) and *Zfp281^f/+^*; MMTV-Cre (cOE) with FVB MMTV-ErbB2/HER2/Neu (MMTV-HER2 onward), an established breast cancer mouse model. The MMTV-HER2 mouse expresses the rat epidermal growth factor receptor 2 in the mammary epithelium upon puberty, causing spontaneous premalignant mammary lesions that lead to primary tumors (PT) after 25 weeks of age **(S. Fig. 1C)**. Early in cancer progression, that is, during the mammary intraepithelial neoplasia state (DCIS in humans), HER2 upregulation leads to early dissemination. DCCs remain dormant for long periods, and metastatic initiation is usually detected at the same time the PT develops.^1,2,21^ Transgenic *Zfp281* cKO (*Zfp281^f/f^*; MMTV-Cre; MMTV-HER2), *Zfp281* cOE (*Zfp281^f/f^*; MMTV-Cre; MMTV-HER2) and *Zfp281* WT (*Zfp281^+/+^*; MMTV-Cre; MMTV-HER2) mice were verified by PCR genotyping **(S. Fig. 1D)**. *Zfp281* WT mice showed expected progression kinetics from our MMTV-HER2 and MMTV-HER2/MMTV-Cre-CFP mice as published.^21^ Specific cKO and cOE of ZFP281 expression in luminal cells of MG was further confirmed by IF staining of MG sections from 10-week-old virgin animals **(Fig. 1C)**. cKO mice showed a complete loss of ZFP281 expression from the luminal cells, while cOE mice showed higher expression than WT. HER2 alone is known to increase branching of the mammary tree.^25^ To examine the dependence of HER2-driven aberrant branching on ZFP281, we analyzed the MG at the early lesion stage in cKO virgin animals. Whole mounts were prepared using inguinal MG from virgin mice (10 weeks old) and post-pregnant mice. ZFP281 cKO animals showed no change in branching caused by HER2.

ZFP281 cOE does not alter mammary tree density compared with WT ZFP281 virgin animals **(Fig. 1D-top panels, Fig. 1E, S. Fig. 1E)**. This argues that additional mechanisms, activated by HER2 and ZFP281-independent, still fuel HER2-driven aberrant branching. We next monitored the effect of pregnancy, a commonly used approach in the MMTV-HER2 model to accelerate progression to early lesions, in WT, cKO, and cOE mice. Interestingly, pregnancy increased expression of ZFP281 in the ELs of WT mice **(Fig. 1F)** and, expectedly, pregnancy enhanced branching of the early-stage MGs regardless of the WT, cKO, or cOE state of the HER2+ luminal cells **(Fig. 1D-bottom panels, Fig. 1E)**. Thus, ZFP281 expression is not essential for the HER2-driven branching program during early lesion phases, and pregnancy signals can override any loss or cOE of ZFP281.

Next, we tested whether ZFP281 cKO and cOE affected mammary cancer initiation and progression. The MMTV-LTR promoter driving HER2 expression is stimulated by hormones during sexual maturity, pregnancy, and lactation.^26,27^ Thus, as described above, mice were bred to expose the mammary tissue to one round of pregnancy and lactation, which potentiates tumorigenesis.^21^ Female mice, ZFP281 WT, cKO, and cOE, that underwent one round of pregnancy, were allowed to live until they developed a PT of 1 cm in diameter. PT data revealed that 95% (20 out of 21 mice) of the WT mice, and 100% (25 out of 25) of the ZFP281 cKO animals developed PT **(Fig. 2A)**. However, by week 40, while ∼80% of WT and cKO animals had developed tumors, 0% of ZFP281 cOE mice had developed overt tumors. In fact, it took another 10 months for the ZFP281 cOE mice to develop tumors, and at a much delayed pace. Only 67% (8 out of 12) of the cOE mice showed PT development at >80 weeks (∼1.7 years) **(Fig. 2A)**. No significant differences were observed in the ages at which PT developed in WT and cKO mice **(Fig. 2B)**. In WT, the median duration until PT development was 30 weeks (95% Confidence Interval = 28-31 weeks, range = 24-48 weeks), and in cKO, the median duration until PT development was 32 weeks (95% CI = 26-34 weeks, range = 24-62). In stark contrast, ZFP281 cOE mice not only reduced tumor formation but also dramatically delayed PT onset, with a median time to PT of 65 weeks (95% CI = 54-71 weeks, range = 42-72 weeks) **(Fig. 2B)**. Impressively, ∼30% of the cOE mice never developed PT until the age of 86 weeks (∼1.8 years), and died due to other non-cancer aging-related illnesses. Thus, ZFP281 is not necessary for HER2-dependent tumor initiation and progression in the primary site. However, when overexpressed, it dramatically interfered with HER2-driven tumorigenesis and, in a significant proportion of animals, conferred lifelong protection against tumor initiation. Although not significant, ZFP281 cKO mice showed a trend toward increased tumor multiplicity, with 1-5 PTs per animal (median = 2). In comparison, WT mice had 1-2 PTs per animal (median = 2), whereas most cOE animals developed only 1 tumor per animal (median = 1) **(Fig. 2C)**. Thus, the frequency of initiating events may be affected but not significantly in the present model. However, animals in cKO with multiple PTs had an onset slightly earlier than those with a lesser number of PTs **(S. Fig. 2A)**. The doubling time (DT) for the PTs to reach the size of 1 cm in WT animals varied widely, ranging from 5 to 30 days (median = 17.28 days, IQR = 20.54 days). On the other hand, the DT in cKO animals was less diverse, with a median DT of 11.64 days (IQR = 7.16 days); the cOE group had a median DT of 21.59 days (IQR = 17.76 days) but still statistically insignificant **(Fig. 2D)**. ZFP281 cOE animals that developed tumors, even after a long delay, had similar DTs as WT but slightly slower than that of the cKO animals. However, phenotypically, the PT in cOE appears different, with a larger nucleus than in WT and cKO **(S. Fig. 2B)**.

**Figure 2.**
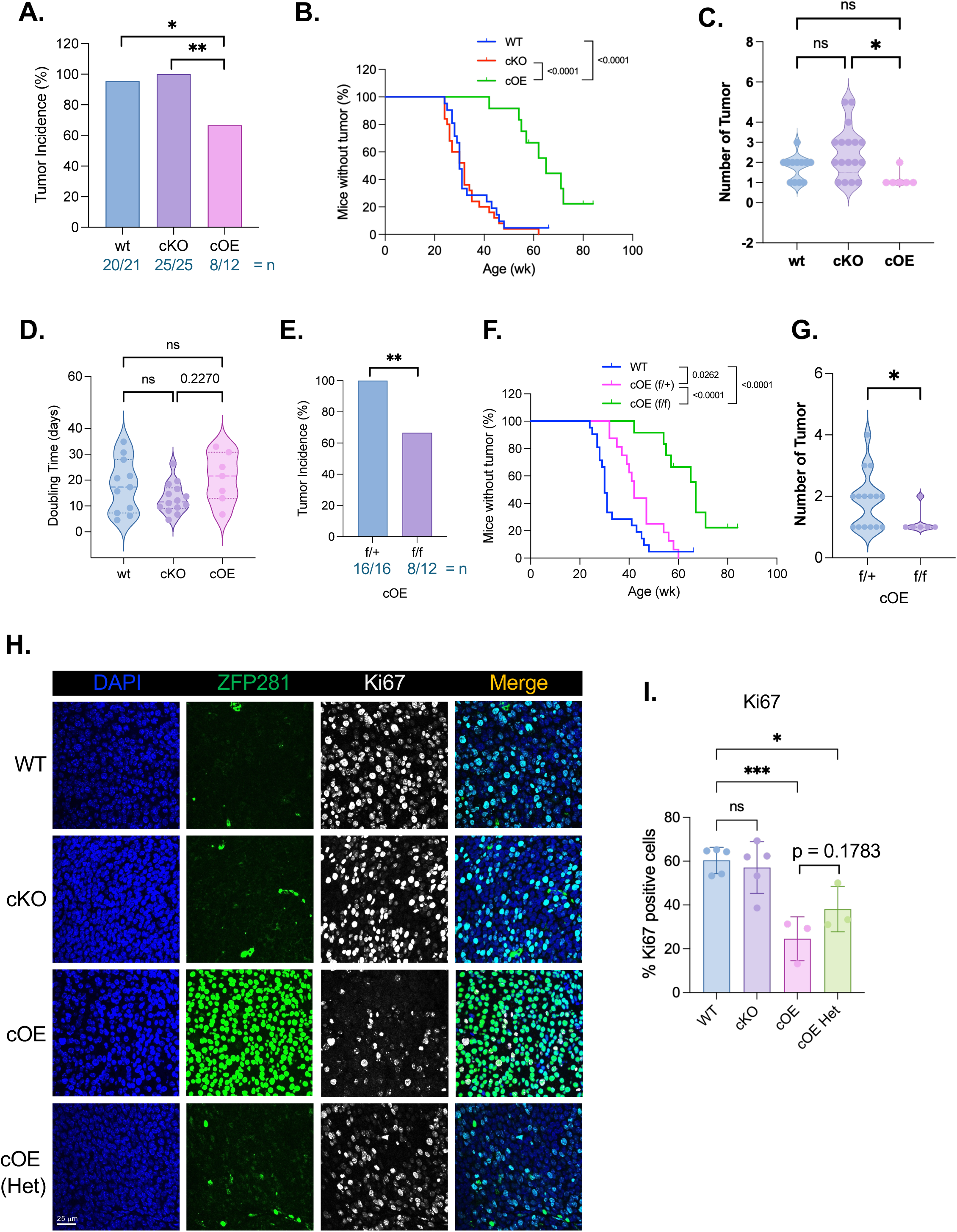
ZFP281 overexpression suppresses primary tumor development. **A.** Percent of mice that developed a primary tumor. Absolute number of animals in each group (number of mice that developed tumors/total number of mice). Statistical test: Fisher’s exact test. **B.** The graph shows the delayed onset of PT development in cOE compared to WT and cKO mice. Animals that did not develop any tumor are marked in vertical small line. Statistical test: Log-rank (Mantel-Cox) test. **C.** The graph shows the total number of tumors developed per mouse until the first PT reached 1 cm in size. Statistical test: one-way Anova (Kruskal-Wallis test). **D.** The graph shows the average doubling time in the PT growth until it reached 1 cm in size. Statistical test: one-way Anova (Kruskal-Wallis test). **E.** Comparison between heterozygous and homozygous OE mice in PT onset. Statistical test: Fisher’s exact test. **F.** Delayed onset of PT development in heterozygous cOE compared to WT but earlier to homozygous cOE mice. Animals that did not develop any tumor are marked in vertical small line. Statistical test: Log-rank (Mantel-Cox) test. **G.** The graph shows the total number of tumors developed per mouse until the first PT reached 1 cm in size. Statistical test: Simple t-test. **H-I.** Immunofluorescence staining of PT for ZFP281 and Ki67 (proliferation marker) and its quantification.

To explore the effect of ZFP281 overexpression and gene dosage, we compared the ZFP281 cOE (heterozygous, F/+) mice with cOE (homozygous, f/f) mice, and interestingly we observed that 100% of F/+ cOE animals developed PT compared to 67% in f/f cOE mice **(Fig. 2E)**. Although F/+ cOE mice developed PT, they developed significantly late (median = 42 weeks, 95% CI = 37-54 weeks, range = 32-60) compared to WT and cKO animals, which developed PT at the median age of 30 and 32 weeks, respectively **(Fig. 2F)**. Thus, even one additional allele of Zfp281 gene dosage (at the H11 locus) can severely impact HER2-driven progression. We also observed increased tumor multiplicity in F/+ cOE mice, ranging from 1-3 tumors per animal (median = 2, IQR = 1) compared to only one tumor per animal in f/f cOE mice **(Fig. 2G)**. Further, we performed IF staining to detect ZFP281 and Ki67 expression in PTs of WT, cKO, and cOE animals **(Fig. 2H)**. We observed a heterogenous population of ZFP281 expressing PT cells in cOE (F/+) animals compared to more consistent expression in cOE (f/f), WT and cKO **(S. Fig. 2C).** As expected, we observed low ZFP281 but high Ki67 expression in WT and cKO PTs **(S. Fig. 2D)**. Interestingly, compared to WT and cKO PT, we did not observe any reduction in ZFP281 expression in homozygous cOE PT but significantly reduced Ki67 expression, suggesting still a measurable inhibitory effect of ZFP281 on proliferation, but also a bypass mechanism (unknown) that allowed for tumor mass expansion in both heterozygous or homozygous overexpression animals **(Fig. 2I, S. Fig. 2C-D)**. Accordingly, we observed a significant decrease in ZFP281 expression in heterozygous OE PTs, comparable to WT and cKO animals, and a decrease in Ki67 expression **(Fig. 2H-I)**. This confirms that ZFP281 cOE suppresses PT development, but loss of ZFP281 does not significantly promote tumorigenicity and tumor multiplicity.

### ZFP281 expression and hormone signaling drive mammary development in virgin mice

Since ZFP281 suppresses PT development in the HER2 mouse model but does not affect branching dynamics under HER2 expression, we wanted to investigate its role in normal MG development. Since the epithelial compartment in the MGs undergoes differentiation during pregnancy, giving rise to milk-producing luminal and contractile basal cells,^28^ and, at the end of pregnancy, undergoes involution through large-scale apoptosis,^28^ it is important to understand ZFP281’s function in MG differentiation, branching, and alveolar development during pregnancy and post-pregnancy. To this end, we utilized ZFP281 WT, cKO, and cOE (f/+) mice from the MMTV-Cre background only (no HER2 oncogene expressed). We performed whole-mount staining of inguinal MG from 10 weeks (virgin), 11-12 weeks (18 days pregnant), and 15-16 weeks old (post-weaned, i.e., 2 days after pups weaning) WT, cKO, and cOE mice **(Fig. 3A and S. Fig. 3A-B)**. Unlike in the HER2+ model, whole-mount analysis showed a significant loss of MG branching in the virgin cKO mice, while cOE animals showed extensive mammary tree branching compared to WT littermates, leading to a higher density of MGs **(Fig. 3A-C)**. This extensive MG branching was similar to the branching in a pregnant female mouse but with underdeveloped terminal end buds (TEBs) **(Fig. 3A)**.^29^ The pregnant and post-weaned mice, however, showed no significant difference in the MG branching upon ZFP281 cKO or cOE **(Fig. 3A and S. Fig. 3A-B)**, suggesting that, as in the early lesion stage, hormonal signaling overrides any loss or gain of ZFP281 gene dosage. Interestingly, MGs in cOE animals were significantly smaller compared to MG in WT or cKO animals **(Fig. 3D)**. However, there was no difference in the size of MG in pregnant and weaned mice (**S. Fig. 3C-D**). Based on these results, we conclude that postnatal MG branching is regulated by ZFP281. However, at least as revealed by whole-tissue mounts, pregnancy signals do not depend entirely on ZFP281 for MG expansion during pregnancy.^21,30,31^

**Figure 3.**
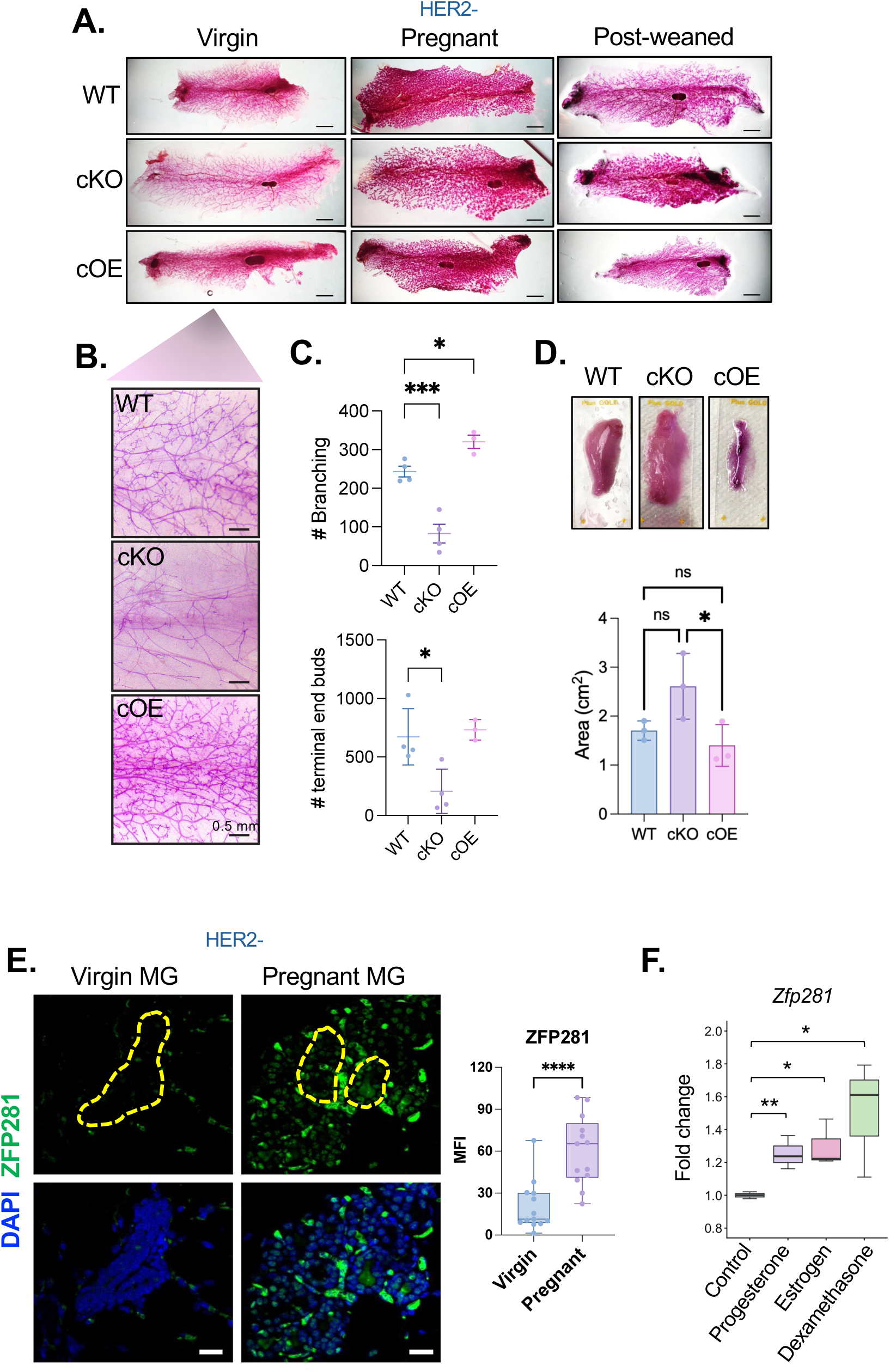
ZFP281 regulates early branching morphogenesis in the normal mammary gland. **A.** Whole mounts of inguinal mammary glands from virgin (10 weeks age), pregnant (10-12 weeks age, 18 days pregnant), and post-weaned (16-18 weeks age, 48 hours after weaning) MMTV-Cre (WT, cKO and cOE) mice. **B-C.** Zoomed-in images of whole mount MG from virgin mice and quantification of the number of branching and terminal end buds. Statistical test: Ordinary one-way ANOVA. **D.** Whole mounts from MMTV-Cre (WT, cKO, and cOE) showing differences in MG size and its quantification. Statistical test: Ordinary one-way ANOVA. **E.** Immunofluorescence staining of ZFP281 in the MG of MMTV-Cre WT virgin and pregnant mice showing induced expression of ZFP281. Yellow boundaries marked individual ducts. The graph shows the quantification. Statistical test: Simple t-test. **F.** The graph shows increased *Zfp281* expression in MMTV-Cre WT (virgin) organoids treated with Progesterone, Estrogen, and Dexamethasone for 6 hours. Statistical test: Ordinary one-way Anova, n=3.

MG development is tightly regulated by hormones secreted during and after pregnancy, especially estrogen (E2), progesterone (P4), and Prolactin.^31–33^ To identify any effect of pregnancy and post-pregnancy on the expression of ZFP281, we conducted IF staining of ZFP281 in the MG sections of virgin and pregnant mice. Pregnant WT mice exhibited a significant increase in ZFP281 expression, indicating a potential regulation of *Zfp281* by pregnancy signals **(Fig. 3E)**. We also noted elevated levels of ZFP281 in post-weaned maternal mice **(S. Fig. 3E)**, suggesting a long-term effect of hormones on ZFP281 expression and likely an important role of ZFP281 in pregnancy. To confirm the elevated expression of ZFP281 due to the pregnancy hormones, we treated ZFP281 WT organoids (*in vitro)* with progesterone (P4, 40 ng/ml), estrogen (E2, 10 nM), and Dexamethasone (a glucocorticoid, 100 nM) for 6 hours and examined the RNA expression by qPCR. P4, E2, and Dexamethasone treatments showed increased expression of *Zfp281*, suggesting a regulatory signal controlling *Zfp281* expression **(Fig. 3F)**. We conclude that in normal mammary glands, ZFP281 is required for post-natal branching morphogenesis. We further conclude that ZFP281 expression is regulated by pregnancy hormones but is dispensable for mammary tree expansion during and after pregnancy. The fact that ZFP281 cOE caused unwanted and uncontrolled branching and that it is upregulated by HER2 signaling in early lesions^21,34^ nevertheless suggested that this function might be co-opted by oncogenes to promote early dissemination.

### ZFP281 maintains hybrid and stem programs in mammary epithelial cells

ZFP281 regulates key pluripotency genes and EMT drivers while promoting a mesenchymal-like dormancy program in early disseminated breast cancer cells.^1,5,8^ To explore how these mechanisms emerged in normal mammary epithelial cells (MECs), we examined the morphology of organoids generated from the MG of ZFP281 WT, cKO, and cOE (f/+) virgin mice (with MMTV-Cre). WT organoids displayed a polarized, epithelial appearance; cKO organoids were larger and resembled morula-stage embryos, while cOE organoids were smaller and more disorganized **(Fig. 4A, Video 1-3)**. Protrusive morphology in the organoids was assessed by calculating the length-to-width ratio, revealing a higher protrusion ratio in organoids from MG of cOE animals, which may be due to the more mesenchymal phenotype^1^ **(Fig. 4B)**. Further, we examined the expression of stem cell, luminal identity, and EMT genes in those organoids by qPCR. We found that ZFP281 cKO led to elevated expression of MET and luminal genes *Klf4* and *Gata3*, while ZFP281 cOE had no significant effect on *Gata3* **(Fig. 4C)**. Interestingly, the regulator *Myc*, involved in the cell cycle, stemness, and metabolism, was upregulated in both perturbations. This likely indicates distinct outcomes in each context that are unrelated to proliferation, since ZFP281 represses cell division while preserving some cell plasticity. Moreover, and in line with a more epithelial-hybrid state, ZFP281 cKO caused an increase in the expression of *Cdh1* (E-Cadherin) and *Zeb1*, but it repressed genes such as *Vim* and *Snail* with no changes in *Cdh2* and *Twist1* (**Fig. 4D**). ZFP281 cOE had a different effect on these genes leading surprisingly to upregulation of *Cdh1,* but expectedly *Vim*, and *Snail* while strongly repressing *Zeb1*, arguing for the cOE tuning a more epithelial-mesenchymal hybrid state in MECs, likely needed during morphogenesis in response to hormonal signals. These findings confirm that ZFP281 levels determine the output of epithelial, hybrid, and mesenchymal gene programs in MECs, with a strong tendency to drive a hybrid-mesenchymal state upon upregulation.

**Figure 4.**
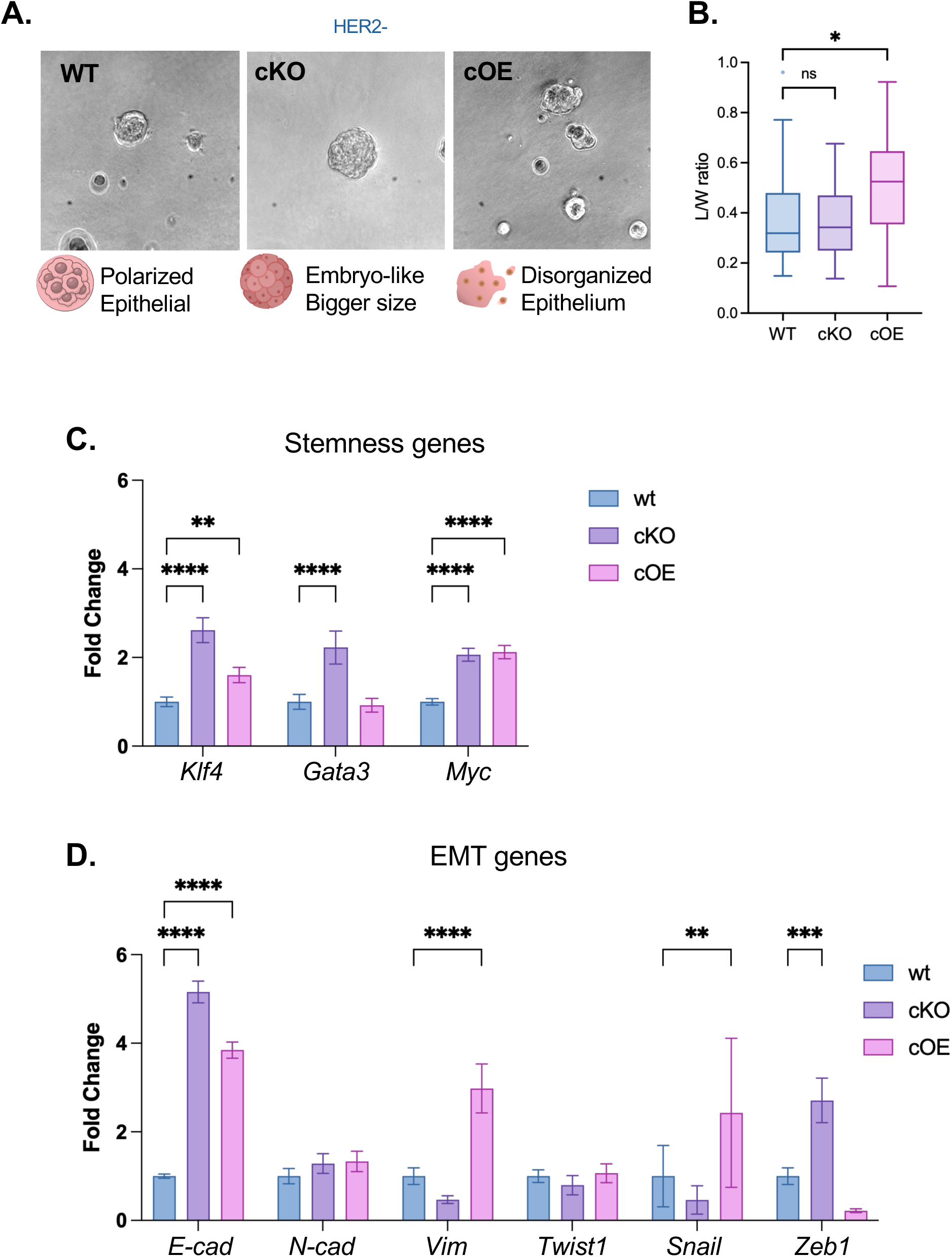
ZFP281 maintains hybrid and stem programs. **A.** Representative images show the phenotype of MMTV-Cre ZFP281 WT, cKO, and cOE mammary gland (virgin mice) organoids. **B.** Quantification showing increased protrusion size (length to width ratio) in cOE organoids compared to WT, reflecting mesenchymal nature. Statistical test: one-way ANOVA. **C-D.** RNA expression of stemness and EMT genes quantified by qPCR in MMTV-Cre ZFP281 WT, cKO, and cOE mammary gland (virgin mice) organoids. Statistical test: one-way ANOVA, n=3.

### ZFP281 cOE enhances early dissemination but provides life-long protection against metastasis

Our previously published^1^ and new results in Figure 3 showed that ZFP281 is upregulated during EL stages. During the EL stages, ZFP281 enables EL cells to disseminate^1^, likely by deregulating the strong branching function it exerts in the postnatal mammary gland (**Figure 3**). We previously showed that when DCCs are lodged in the lung, TGFβ signaling from alveolar macrophages maintains ZFP281 upregulation and keeps DCCs in a dormant state for extended periods.^2^ This argues that if ZFP281 expression and function are maintained at high levels, metastasis could be blocked by rendering DCCs indolent. To test this possibility, we investigated DCC seeding in the lungs of MMTV-HER2 ZFP281 -WT, -cKO, and -cOE mice at the EL stage, when PTs have not yet developed.^21^ Loss of ZFP281 in cKO mice did not impair dissemination compared with WT animals, suggesting that ZFP281 function can be replaced by other mechanisms in its absence. However, we observed a significantly higher number of DCCs in the lungs of cOE animals compared to WT and cKO mice, further supporting our previous findings of increased dissemination associated with elevated ZFP281 expression **(Fig. 5A-B)**.

**Figure 5.**
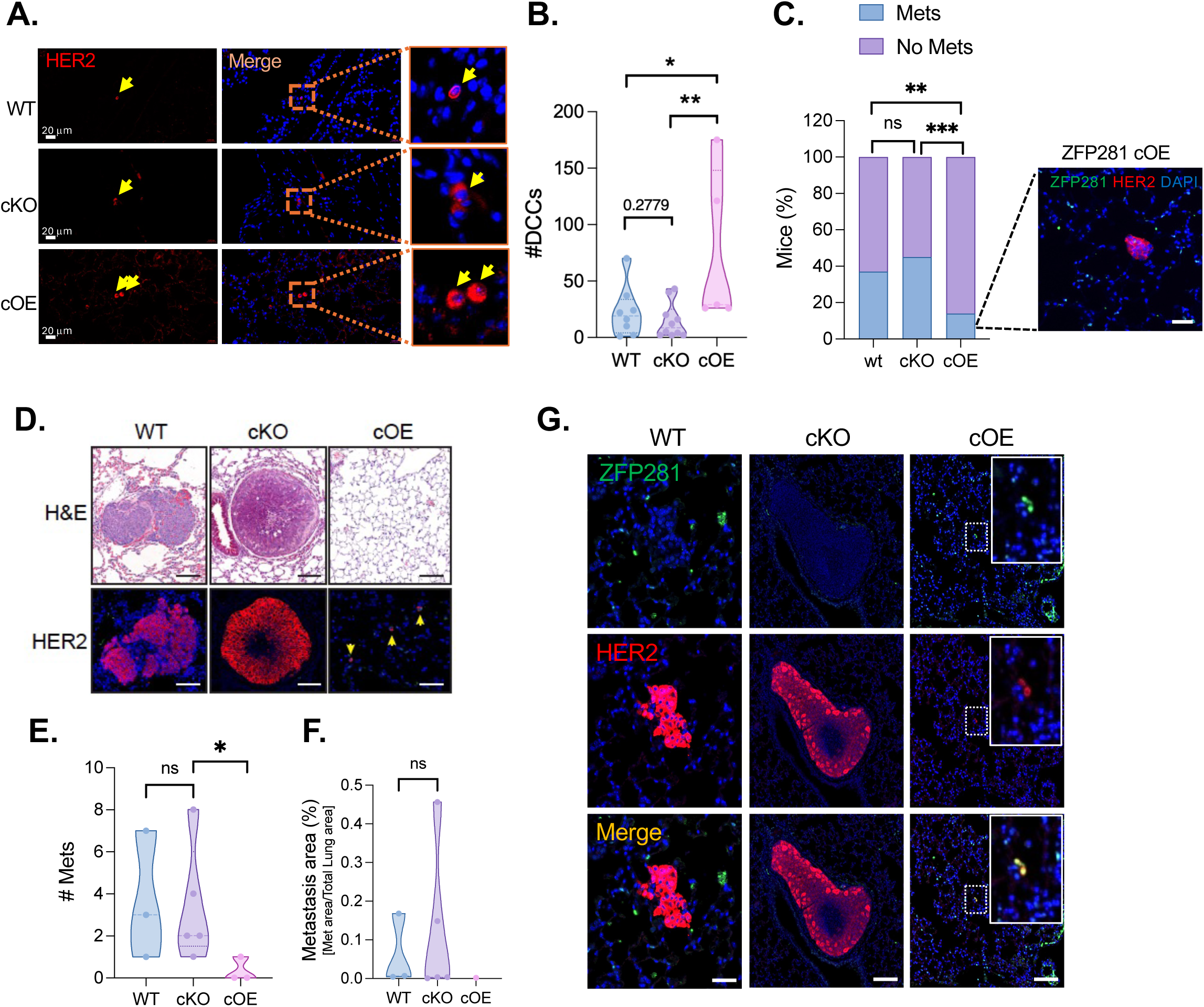
ZFP281 overexpression suppresses metastasis. **A.** HER2 staining of lung sections from post-pregnant MMTV-HER2 ZFP281 WT, cKO, and cOE mice showing DCCs in the lungs. Zoomed images are on the extreme right, and yellow arrows point to DCCs. **B.** The graph shows the quantification of DCCs in the lungs. Statistical test: Mann-Whitney test, n ≥ 5. **C.** Percent of mice that developed metastasis. The inset shows a micrometastatic lesion in a cOE mouse. Statistical test: Post hoc Fisher’s exact tests, n ≥ 5. **D.** Representative images show metastasis in the lungs of ZFP281 WT and cKO mice by H & E staining and HER2 immunofluorescent staining. Single DCCs (yellow arrows) in cOE lungs were visualized by HER2 staining. **E-F.** Number of metastatic lesions per mouse and the percent of total metastatic area normalized to total lung area. Statistical test: Mann-Whitney test, n ≥ 3. **G.** Representative IF images show no ZFP281 expression in the metastatic lesions in the lungs of WT and cKO mice, and ZFP281 expression in the single DCCs in the lungs of cOE mice.

Importantly, when we examined the lungs of MMTV-HER2 ZFP281-WT, -cKO, and -cOE mice that developed PT (or not in the case of ZFP281 cOE), 40% of WT animals developed metastatic lesions, increasing to ∼50% in the absence of ZFP281 in the cKO mice. However, only 1 mouse (10%) in the cOE group showed a small micro-metastatic lesion despite having a > 2-fold increase in solitary DCC burden **(Fig. 5C)**. Both WT and cKO animals showed large metastatic lesions, which showed a more epithelial morphology in cKO animals **(Fig. 5D),** suggesting an enhancement of metastasis upon ZFP281 loss. In stark contrast, the cOE animals showed single, solitary, or small clusters of DCCs throughout the lung (visible only with HER2 staining, not H&E) **(Fig. 5D),** and, in only one mouse, a rare micrometastasis **(Fig. 5C inset)**. The frequency of macro-metastatic lesions in WT and cKO mice lungs ranged from 1-7 to 1-8 mets, respectively. On the other hand, the majority of cOE mice lungs showed no macro-metastasis **(Fig. 5E)**. Similar to the frequency of metastatic lesions in WT and cKO mice, there was no significant difference in the total metastatic burden (met area normalized to total lung area) **(Fig. 5F)**. This argues that while ZFP281 promotes the dissemination of HER2+ cells, it is dispensable for dissemination, but it locks the DCCs in a dormant state for almost the entire life of the mouse after EL development. Critically, even when ZFP281-suppressive mechanisms were bypassed, and tumors developed, metastasis remained suppressed. This reinforces the idea that ZFP281 expression is a powerful bottleneck for metastasis, making its loss necessary to awaken. As reported^1^ and expected, we observed no ZFP281 expression in metastatic lesions in the lungs of WT and cKO animals. By contrast, solitary dormant DCCs in the cOE lungs expressed ZFP281 **(Fig. 5G)**. These observations suggest that continued expression of ZFP281 in DCCs kept them dormant and non-proliferative in cOE animals.

### ZFP281 orchestrates mammary transcriptional programs in a context-dependent manner

To better understand the gene expression program and pathways regulated by ZFP281, and how these compare with those in normal MECs, we conducted RNA-sequencing (RNA-seq) analysis of MG from MMTV-HER2 ZFP281-WT, -cKO, and -cOE mice under virgin and post-pregnancy/EL conditions. In virgin animals, we performed RNA-seq on mammospheres derived from the MG and cultured for 7 days. Principal component analysis (PCA) plot showed that, in the virgin state, the WT and cKO groups cluster together, whereas the cOE group is distinctly separated **(S. Fig. 4A)**. Note that our RNA-seq data confirmed a significant increase of *Zfp281* expression in cOE animals **(S. Fig. 4B)**. In virgin mice, the absence of ZFP281 affected only a small number of genes compared to WT controls, with 20 downregulated and one upregulated **(Fig. 6A, S. Fig. 4C, S. File 1)**. Given that, in its virgin state, the basal expression level of ZFP281 is already considerably low **(Fig. 1C and S. Fig. 4B)**, we expected that its depletion would affect only a limited number of genes. This is also consistent with the fact that ZFP281 cKO did not affect MG branching in the MMTV-HER2 background **(Fig. 1D-E)**. However, KEGG pathway analysis of the downregulated genes in virgin ZFP281 cKO MGs showed enrichment for HIF1α signaling, the Pentose phosphate pathway, and fructose and mannose metabolism, suggesting potential regulation by ZFP281 **(Fig. 6B)**. This also suggests that ZFP281 may function upstream of, or in coordination with, HIF1 to maintain a quiescent state.^35^ Overexpression of ZFP281 resulted in the upregulation of 80 genes and the downregulation of 38 genes **(Fig. 6C, S. Fig. 4C, S. File 1)**. Upregulated genes showed enrichment for glutathione metabolism and cytochrome-mediated xenobiotic metabolism (*Gsta3, Gsta2, Gstt1*), whereas downregulated genes were enriched for ECM-receptor interaction (*Cola1, Spp1, Dmp1*), the AGE-RAGE signaling pathway (*Egr1, Serpine1*), and the IL-17 signaling pathway (*Fos, Ptgs2*) **(Fig. 6B, S. File 1)**. The latter suggests that ZFP281 suppresses the IL-17-driven inflammatory signature and interferes with ECM crosstalk.

**Figure 6.**
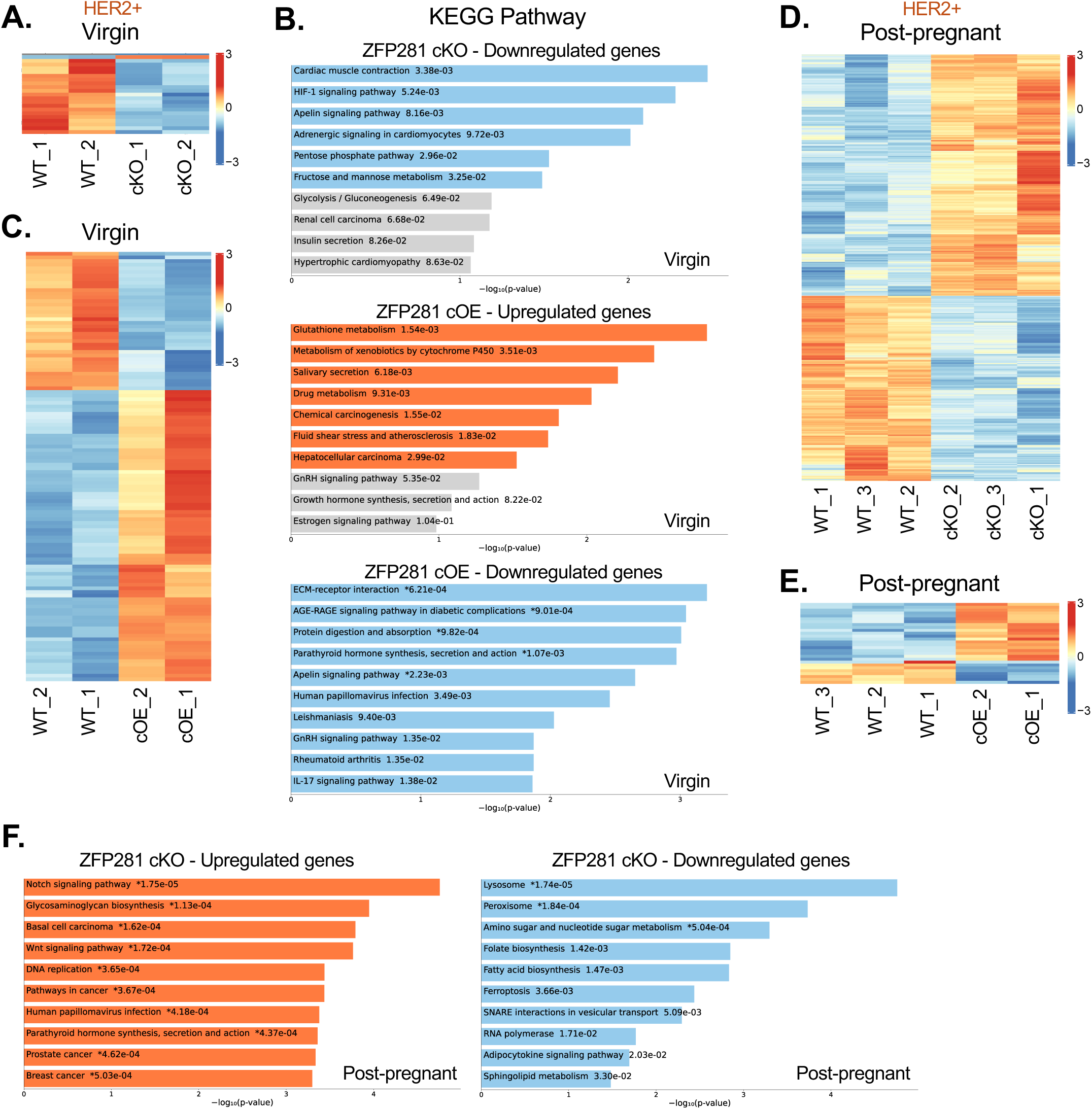
ZFP281-regulated gene expression. **A-C.** Bulk RNA-sequencing analysis in the MMTV-HER2 ZFP281 WT, cKO, and cOE mammospheres prepared from virgin (10-week-old). *A and C* show the genes significantly differentially expressed. *B* shows the KEGG pathway enrichment analysis, highlighting pathways enriched for the significantly upregulated and downregulated genes. **D-E.** Bulk RNA-sequencing analysis in the MMTV-HER2 ZFP281 WT, cKO, and cOE organoids prepared from post-pregnant (16-18-week-old). **F.** KEGG pathway enrichment analysis, highlighting pathways enriched for the significantly upregulated and downregulated genes. Genes significantly upregulated and their associated pathways are shown in the orange bar, and genes significantly downregulated, and their associated pathways are shown in the blue bar. Insignificant pathways are in grey color bars.

In the post-pregnant condition, PCA clustering was different; the WT and cOE groups cluster together, whereas the cKO group formed a separate cluster **(S. Fig. 4D)**. The clustering **(S. Fig. 4A and D)** is consistent with the observation that pregnancy upregulates the expression of ZFP281 **(Fig. 1F and S**. **Fig. 4E, S. File 1),** bringing the WT and cOE samples together. Compared to WT controls, loss of ZFP281 in the EL state leads to differential expression of 2282 genes (1292 upregulated and 990 downregulated), whereas ZFP281 cOE affected only 28 genes (20 upregulated and 8 downregulated) **(Fig. 6D-E, S. Fig. 4F)**. This is consistent with ZFP281 expression observed in our RNA-seq samples, high ZFP281 in WT animals (induced by pregnancy hormones) and cOE did not further increase ZFP281 expression **(S. Fig 4E)**. Upregulated genes upon ZFP281 loss show enrichment for Notch signaling (*Notch1, Jag1, Jag2, Dll1*), Wnt signaling (*Wnt10b, Wnt5a, Wnt5b, Ror2*), and DNA replication pathways (*Mcm2,3,4, Lig1, Fen1*), whereas downregulated genes show enrichment for Lysosome (*Ctsa, Cd63, Atp6v0b*), Peroxisome (*Acot8, Acsl1, Mpv17l*), Ferroptosis (*Prnp, Acsl1, Fth1*), and SNARE interactions in vesicular transport pathways (*Stx16, Stx3, Vamp4*) **(Fig. 6F, S. File 1))**. However, we did not observe KEGG pathway enrichment in either upregulated or downregulated genes upon ZFP281 cOE. Overall, KEGG pathway analysis in post-pregnant animals suggests that ZFP281 inhibits proliferative or oncogenic programs by suppressing Notch and Wnt signaling. Since ZFP281 promotes a mesenchymal-like dormancy gene program during the EL stage and early dissemination, we analyzed the post-pregnant RNA-seq data for gene signatures representing dormancy, EMT, Ep-like, and M-like, curated from Nobre *et al.*^1^ Our analysis revealed that mesenchymal genes such as *Snai1, Twist1*, and *Vim* were upregulated only in WT and ZFP281 cOE MG but not in ZFP281 cKO, whereas *Cdh1* and *Foxa1* were downregulated in the presence of ZFP281, confirming the requirement of ZFP281 for the mesenchymal-like gene signature **(S. Fig. 4G)**.

### ZFP281-linked gene signatures associate with relapse dynamics in DCIS and IBC patients

Collectively, our data suggest that ZFP281-regulated genes suppress growth programs, thereby leading to a long-lived dormant state as long as ZFP281 expression persists. We hypothesized that detecting M-like and Ep-like DCC signatures^1^, enriched for ZFP281 targets, in human samples might identify lesions that preferentially seed dormancy-prone or pro-awakening DCCs. We projected these signatures onto the METABRIC (Molecular Taxonomy of Breast Cancer International Consortium) integrative clusters (ICs) defined by Rueda *et al.*^36^, assuming that transcriptional programs in profiled primary tumors reflect DCC states are largely retained at distant sites. Because we lack information on unsampled precursor lesions, we restrict our interpretation to relapse driven predominantly by DCCs disseminating from the resected primaries.

Across ER+/HER2-ICs, most subtypes were Ep-skewed, whereas IC3, IC4ER+ and IC4ER- were clearly M-shifted. Late-recurring high-risk clusters (IC1/2/6/9) were uniformly Ep-dominant, while M-dominant IC3 and IC4ER+ fell into a lower-risk group, consistent with M-dominance tracking with reduced awakening **(Fig. 7A)**. Late-risk ER+ ICs displayed strong Ep-like medians with M-like scores near or below zero but broad upper tails of M-high tumors, suggesting that predominantly Ep-like primaries harbor sub-clonal or patient-specific M-like/dormant DCCs that may later revert to a proliferative Ep-like state, fitting their slowly rising late-relapse curves. Lower-risk, strongly M-shifted ER+ ICs (IC3, IC4ER+) showed substantially lower 20-year distant relapse/death probabilities, supporting the idea that stable M-like programs (such as those enforced by ZFP281 overexpression^1^ and in this paper) have limited plasticity to switch toward a highly Ep-like proliferative state, thereby seeding DCCs with a lower probability to switch to an aggressive Ep-like outgrowth. The latter finding is consistent with the association of persistent M-like/ZFP281+ states with long-term dormancy and the requirement for loss of this program plus Ep acquisition to drive lethal metastasis. Good-prognosis IC7/8, with both signatures near zero, exhibited a low late relapse risk, consistent with a homogeneous, less plastic population that rarely enters either a strongly M-like dormant state or an extreme Ep-like proliferative state. TNBC (triple negative breast cancer) IC10 was strongly Ep-skewed, with high early recurrence but low late recurrence risk, consistent with a scenario in which DCCs fail to engage durable M-like dormancy. These patterns show that the most informative signal is not whether an IntClust is globally M- or Ep-dominant, but the distribution within the cluster, especially the upper tail of M-like scores in Ep-dominated late-risk groups. Late-recurring ER+ ICs are Ep dominant yet heterogeneous in M-like scores, while some lower-risk ER+ ICs are more consistently M-shifted but have better long-term outcomes, a pattern consistent with a ZFP281/M-like dormancy model.

**Figure 7.**
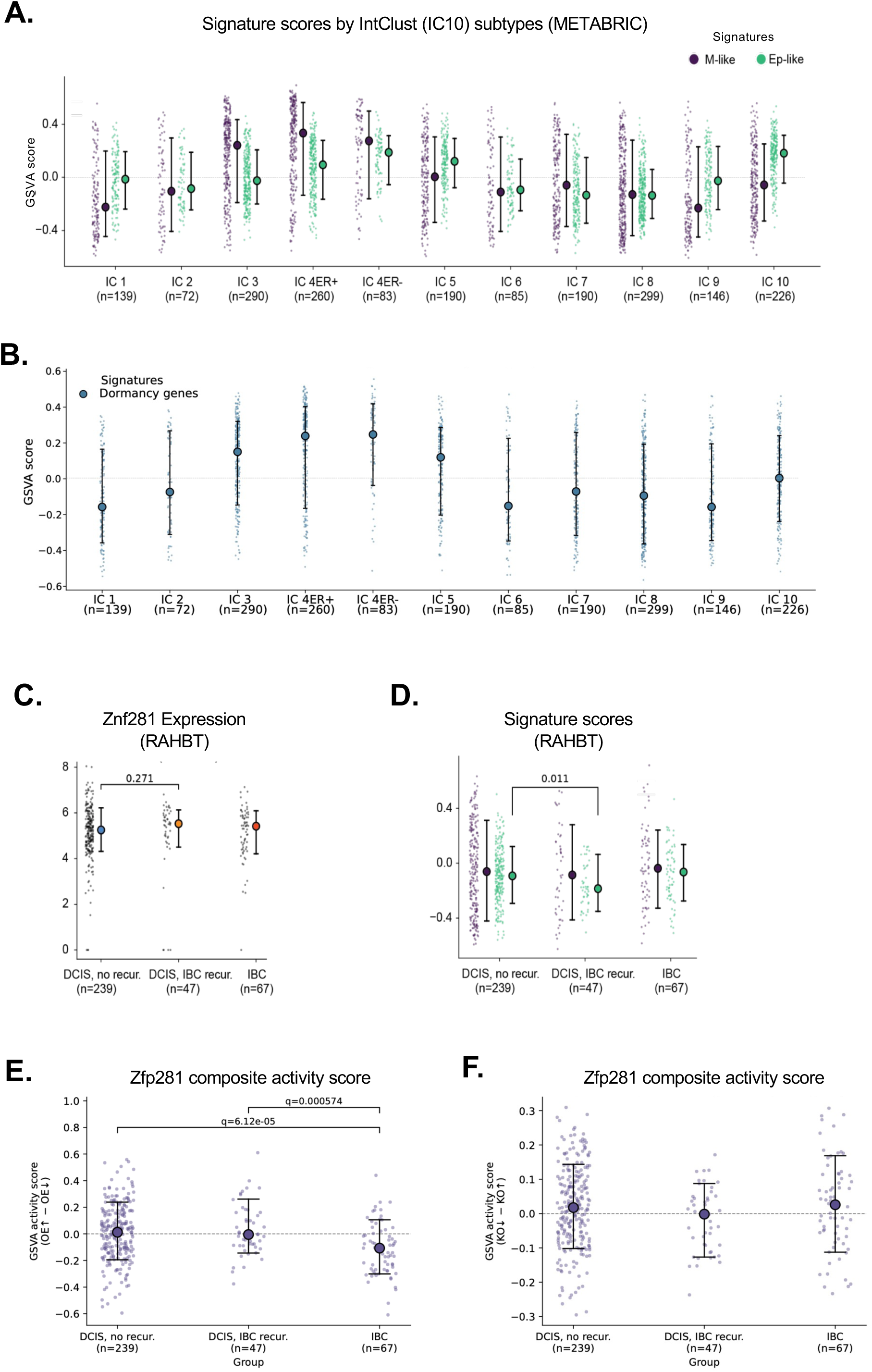
ZNF281 (ZFP281)-linked gene signatures apprise relapse dynamics in breast cancer patients. **A.** GSVA (Gene Set Variation Analysis) score of ZFP281-linked M-like and Ep-like gene signatures from Nobre et al.^1^ in the METABRIC IntClust (IC10) molecular subtypes. IC3, IC4ER+, IC7 and IC8 are low-risk ER+ subtypes; IC10 and IC4ER- are TNBC subtypes having variable relapse patterns; IC1, IC2, IC6 and IC9 are high-risk late-relapsing ER+ subtypes; and IC5 is a HER2+ subtype. Each point represents one tumor sample; large circles indicate median values; vertical lines span the 15th to 85th percentiles. The number of patients in each subtype is shown in parentheses. **B.** GSVA score of a dormancy signature^1^ in the IC10 subtypes. **C.** *Znf281* RNA expression in the primary tumor specimens in breast cancer patients with DCIS (ductal in situ carcinoma) with and without recurrence, and IBC (invasive breast cancer) using the Laser Capture Microdissection (LCS) from the Resource of Archival Breast Tissue (RAHBT). n = number of patients is in parentheses. **D.** GSVA score of M-like (purple) and Ep-like (green) gene signatures in DCIS patients (with and without recurrence) and IBC (Mann–Whitney test, p = 0.01). **E.** Zfp281 composite (upregulated and downregulated genes upon ZFP281 OE in virgin animals) activity score; and **F.** Zfp281 composite (upregulated and downregulated genes upon ZFP281 KO in post-pregnant animals) activity score in DCIS with and without recurrence, and IBC.

We next projected the dormancy (D) signature score^1^ onto METABRIC ICs **(Fig. 7B)**. Low-risk IC3 and IC4ER+ exhibited positive D-score medians comparable to their M-like scores, indicating enrichment for both M-shifted and dormancy programs, whereas good-prognosis IC7/8 had M>E but D-score medians near zero with narrow dispersion. High-risk late-recurring ER+ clusters (IC1/2/6/9) showed D-score medians near or below zero, confirming these groups lack a strong dormancy signal at the IC-median level. However, these groups still have high D-score events that, like M-like scores, may identify events that switch later from a dormant to a proliferative state. In TNBC, IC4ER- showed an M-shifted, D-score-high profile, while IC10 was Ep-skewed with D-score medians around zero, mirroring their clinical behavior as a dormancy-enriched, late-risk subgroup versus an early-relapsing, largely non-dormant TNBC group. Together, these data indicate that the D-score provides information concordant with, but not redundant to, the M/Ep signatures.

To gain insight into early dissemination and the contribution of M- and Ep-like signatures, we also analyzed M-like and Ep-like signature scores in ∼100 laser-capture microdissected DCIS specimens that either did not recur or were associated with IBC (invasive breast cancer) local recurrence. ZNF281 expression trended non-significantly to higher levels in samples that recurred as IBC **(Fig. 7C)**. However, Ep-like scores were significantly lower in DCIS with recurrence versus no recurrence (Mann–Whitney test, P = 0.011) **(Fig. 7D)**. These findings suggest that DCIS lesions with a higher M/E ratio are enriched for transcriptional programs associated with early dissemination and dormancy, and are more likely to recur as IBC, while lesions with balanced M=E signatures may lack the M-like program associated with local dissemination. Because we do not have information on distant relapse, we can only interpret the signatures’ contribution to local dissemination and recurrence.

We next explored ZFP281 signatures **(S. File 2)** derived from virgin HER2+ early lesions (WT, cKO, and cOE) in the DCIS cohort. The composite ZFP281 virgin signature showed a graded decrease from DCIS to IBC, with IBC displaying significantly lower ZFP281 activity than either non-recurrent or IBC-recurrent DCIS **(Fig. 7E)**, consistent with the loss of a ZFP281-associated dormancy program during progression, as also detected at the protein level in human DCIS vs IBC.^1^ Within DCIS, however, the composite score did not distinguish lesions that later recurred from those that remained non-recurrent. Notably, only genes repressed by ZFP281 overexpression (cOE-down) showed a robust shift between DCIS and IBC **(S. Fig. 5A),** suggesting that de-repression of these ZFP281-suppressed targets is the most informative feature of the virgin signature in the context of progression. The ZFP281 cOE-down signature includes inflammatory and IL-6 family cytokine signaling genes *(Il11, Ptgs2, Chil3*), growth factor and angiogenic regulators, and a prominent ECM/remodeling and stiffness program (*Spp1/Opn, Postn, Serpine1, Col1a1*) as well as genes linked to enhanced adhesion, perivascular/niche interactions, and myeloid or stromal crosstalk **(S. File 2)**. Thus, loss of ZFP281 during progression would derepress an inflammatory, ECM-activated, and angiogenesis-permissive state. Using signatures derived from virgin and post-pregnancy mouse models, the composite ZFP281 activity score did not discriminate DCIS that did and did not recur **(Fig. 7F, S. Fig. 5B)**. However, the cKO-down signature showed that DCIS that did not recur retained higher expression of genes that require ZFP281 for induction, whereas DCIS that later progressed to IBC showed a significant reduction in this cKO-down score. This cKO-down module comprises a broad stress-adaptation and dormancy-supportive network, including redox and metabolic buffering genes, autophagy/lysosome and vesicle trafficking, quiescence and growth-restraint regulators, as well as immune-modulatory or stromal factors **(S. File 2)**. Together, these data suggest that the gene signature derived from the post-pregnancy cKO-down (ZFP281-induced) informs on the stress-autophagy-quiescence program that distinguishes DCIS lesions that later recur. However, whether this module is linked to parity in human DCIS remains to be determined. The direction of this association, loss of ZFP281-induced genes tracking with progression, is consistent with the dormancy-gatekeeper role observed in mice.

## DISCUSSION

Previously, we demonstrated that downstream of HER2, ZFP281 upregulation activates an M-like program in DCCs, which is associated with EMT and primed pluripotency programs in early mammary cancer lesions. ZFP281 upregulation permits early cancer cells to disseminate while concurrently enforcing a dormancy program upon arrival to target organs, which is further enforced by the lung niche.^2^ We proposed^1^ that this mechanism of ZFP281-driven early dissemination may co-opt programs activated during mammary gland branching morphogenesis and that ZFP281 functions as a gatekeeper of dormancy in DCCs post-extravasation. Here, we discuss key findings of our results that support and expand the discovered mechanisms and their potential impact on human breast cancer.

Our data from studies of normal mammary gland development in virgin cKO animals clearly show that ZFP281 is required for mammary tree branching. This was confirmed by our finding that ZFP281 cOE causes what appeared to be relentless branching, which also limits duct expansion due to frequent branching. This excessive ductal side branching may now allow sufficient time for MECs to differentiate. We observed a reduction in *Ror2* (Wnt signaling) and *Notch 1* and *Notch 2* (Notch signaling) in ZFP281 cOE virgin animals, which were previously shown to be required for controlled MG branching morphogenesis.^37,38^ Interestingly, during pregnancy, ZFP281 was upregulated in luminal and basal cells, and its expression in organoids can be stimulated by pregnancy hormones; the link to pregnancy and branching programs was also seen in the PCA analysis, where RNA-seq profiles of WT HER2+ clustered close to ZFP281 cOE HER2+ cells, which was not seen in RNA-seq samples from virgin animals. However, during pregnancy, branching proceeded in an apparently normal fashion in conditional ZFP281 KO or OE mice. Thus, while mammary tree morphogenesis in virgin animals requires ZFP281, pregnancy hormones tap into redundant mechanisms that override the need for ZFP281 in branching and alveologenesis, likely ensuring the preparation of the MG for lactation.^28,33^ These data confirm our initial prediction that ZFP281 is required for initial MG branching, and that the HER2 oncogene co-opts its function, enabling early, unwanted dissemination.

Our data support that loss of ZFP281 does not accelerate HER2 primary tumor formation, suggesting that it does not function as a typical tumor suppressor.^39^ However, cOE of ZFP281 markedly constrained PT formation, suggesting that it must be downregulated, not deleted, at some point to favor HER2-driven growth programs. ZFP281 enforced expression had such a dominant function that HER2+ animals either never developed PTs or did so very late, and the median age was extended to 59 weeks, compared to 30 weeks in cKO and WT animals, respectively. Considering that mice live ∼100-130 weeks, ZFP281 overexpression protected mice from developing HER2 tumors for the equivalent of a quarter of their life. We interpret that cOE of ZFP281 can override HER2’s transformative effects by constraining its transformation and plasticity programs to an M-like state that prevents active proliferation. This supports the idea that, in advanced PT lesions in mice and humans,^1^ ZFP281 is downregulated to a threshold that allows epithelial programs to take over, coincident with primary tumor expansion.^40–42^ This gene/protein dosage hypothesis was supported by our analysis of PT development in ZFP281 cOE heterozygous animals.

Does enforced ZFP281 expression produce the same effects on metastatic outgrowth? As our previous findings predicted,^1^ ZFP281 cOE significantly enhances dissemination to the lungs. This dataset also indicates that recruitment of the ZFP281 program by HER2 activity occurs very early, locking early-lesion cells in a growth-arrested yet highly disseminating M-like gene program. This program must be suppressed for growth to proceed, as shown by the low ZFP281 expression in mouse and human IBC and in mouse overt metastatic lesions. What we did not expect was the remarkable robustness of ZFP281 cOE’s pro-dormancy and anti-metastatic effects in HER2+ DCCs. While 40-50% of ZFP281 WT and cKO animals developed large lung metastases, and DCCs increased in ZFP281 lungs by about six-fold, these DCCs remained in a dormant single-cell state. This was not a short-term phenomenon; instead, it was a lifelong trait in ZFP281 cOE DCCs. Single DCCs in the cOE animal’s lung showed elevated and sustained ZFP281 expression, and only one animal developed a micro-metastasis. The increased number of DCCs in the lungs of ZFP281 cOE mice was significant because it ruled out the possibility that metastatic suppression depended on reduced primary tumor growth. Therefore, if ZFP281 is enforced in DCCs at secondary organs, it can have a lifelong impact on preventing metastasis.

In the METABRIC IBC cohort analysis, the balance of M-like versus Ep-like signatures and the dormancy score showed that ER+ ICs with Ep-dominant profiles and low D-scores (e.g., IC1/2/6/9) experienced earlier relapse and cancer-specific death, while lower-risk ER+ clusters with stronger M-like and dormancy signatures (IC3, IC4ER+, IC7/8) had significantly delayed relapse, often beyond 20 years.^1,2,36,43^ These patterns suggest that ZFP281-linked M-like and dormancy programs, similar to those in the conditional mouse models, influence both lesion progression and the timing of DCC reactivation, reinforcing the idea that ZFP281 functions as a dormancy gatekeeper. Its erosion shifts human breast cancers toward earlier metastatic outgrowth.

In human DCIS, ZFP281-associated programs influenced relapse risk in a stage- and context-dependent manner. Loss of the ZFP281-repressed OE-down module (from virgin HER2+ mammary glands), enriched for inflammatory, angiogenic, and ECM-remodeling genes, marked progression from DCIS to IBC, but did not distinguish recurrent from non-recurrent DCIS. In contrast, the significant repression of Ep-like scores in recurrent DCIS, combined with trending higher ZNF281 expression, supports a model in which DCIS lesions prone to local invasive relapse acquire an M-biased phenotype, consistent with clinical evidence that a minority of DCIS contain cells capable of local dissemination, long-term dormancy, and late recurrence.^44–46^

Importantly, both the virgin and post-pregnancy composite ZFP281 activity scores failed to stratify DCIS relapse risk as groups, although the graded decrease from DCIS to IBC in the virgin signature confirms that erosion of ZFP281-dependent programs accompanies overt progression. By contrast, using signatures derived from the post-pregnancy mouse model, DCIS with subsequent IBC recurrence were selectively depleted of the ZFP281-induced “KO-down” module, a stress-adaptation and dormancy-supportive program, whereas non-recurrent DCIS retained higher KO-down scores. These patterns are consistent with ZFP281 acting as a dormancy gatekeeper whose loss shifts human breast cancers toward earlier metastatic outgrowth, and the M-like/Ep-like and dormancy signatures may offer more informative stratification of relapse dynamics across breast cancer subtypes.^1,2^

Mechanistic insights from the RNA-seq analysis reveal that ZFP281 cOE cells may coordinate HIF1α transcription, ferroptosis, and fatty acid biosynthesis pathways, while also inducing glutathione and xenobiotic metabolism. Recently, it was shown that dormant cancer cells in response to HER2 signaling inhibition upregulate redox detoxification pathways^47^ and that dormant mammary cancer DCCs upregulate de novo lipogenesis and favor the incorporation of monounsaturated fatty acids (MUFAs) into their cellular membranes through the activation of acyl-coenzyme A synthetase long-chain family member 3, which limits ferroptosis.^48^ These pathways may represent potential vulnerabilities for fully eradicating ZFP281+ dormant DCCs. Approaches that could induce and maintain ZFP281 programs may be therapeutically exploited to eliminate the capacity for distant recurrence in breast cancer cells. This, combined with drugs that target the vulnerabilities of these ZFP281+ dormant DCCs, may enable the eradication of stably dormant cancer cells.

## METHODS

### Mouse models

The method used to generate mouse models is briefly described in the results section. Detailed methods and strategies employed to create these mouse models are provided in the **supplementary information**. All mouse experiments were conducted in accordance with the guidelines approved by the Institutional Animal Care and Use Committee (IACUC) at Albert Einstein College of Medicine and Columbia University Irving Medical Center. Tumors were not allowed to grow beyond the IACUC-allowed limit of 1 cm in length.

### Mice experiments

Mice were euthanized using isoflurane and cervical dislocation. All 5 pairs of mammary glands were examined for visible lesions or palpable tumors. Mammary glands were collected and processed to generate mammospheres and organoids as described below. Mice were perfused with PBS, and organs were collected. For histopathology, organs were fixed in 4% paraformaldehyde (PFA, Thermo Scientific, 28908) for 48 hours, processed, embedded in paraffin, and sectioned.

### Whole Mount staining

The fourth inguinal MG was removed and directly spread on the glass slide. The tissue was fixed in the formalin solution overnight, followed by fixation in Carnoy’s solution (a mixture of ethanol, chloroform, and glacial acetic acid in a 6:3:1 ratio) for 4 hours at room temperature. The tissue was gradually hydrated in a decreasing ethanol gradient and stained with alum carmine solution for at least 24 hours. The tissue was dehydrated in an increasing ethanol gradient and cleared in Xylene overnight. Stained whole-mount MG was stored in methyl salicylate, and images were captured using a Nikon SMZ800 stereo microscope.

### Mammosphere and organoid culture

Whole mammary glands were minced with a sharp blade and then digested in 0.15% Collagenase 1 A (Sigma, C-9891), 2.5% bovine serum albumin (BSA, Fisher Scientific, BNP1605) at 37 °C with agitation for 30 min. Red blood cell lysis buffer (eBioscience, 4333-57) was used for 2–5 min, and cells were filtered through a 40-μm filter. For mammosphere culture, 5 × 10^5^ cells were cultured in mammosphere medium (DMEM-F12, Gibco, 11320-033; B27 supplement, Gibco, 17504-044; Epidermal Growth Factor, Peprotech, AF-100-15-A). For organoids, 5 x 10^5^ cells were cultured in organoid medium (advanced DMEM, Gibco, 12634028; 5% FBS, Gemini, 900-108-500; 10 ng/ml EGF; 20 ng/ml Fibroblast Growth Factor 2, R&D Systems, 233FB025; 4 mg/ml Heparin, Sigma, H3149; 5 mM ROCK inhibitor Y-27632, StemCell, 50-197-5898; and 4% Matrigel, Corning, CB-40230C). Live cell imaging of organoids was performed using a Nikon W1 Spinning Disk confocal microscope.

### Immunofluorescence

Paraffin-embedded tissue sections were hydrated in xylene (twice for 10 minutes), then rehydrated in graded ethanol (3 minutes each), and finally washed with water for 5 min twice. Antigen retrieval was performed in 10 mM citrate buffer, pH 6.0 (Na_3_H_6_H_5_O_7_), for 40 minutes in a steamer. Blocking was performed with 3% BSA in PBS containing 5% normal goat serum (Thermo Fisher, PCN5000) for 1 hour. Antibodies used are summarized in the **supplementary information-Table 1**. Tissue sections were washed with PBS three times and incubated with Alexa Fluor conjugated secondary antibodies (Invitrogen, 1:1000 dilution) at room temperature for 1-2 hours in the dark. Tissues were washed twice with PBS, followed by background autofluorescence quenching using an autofluorescence quenching kit (Vector, NC2046752). Tissues were washed twice with PBS. For ZFP281 detection, the Alexa Fluor™ 488 Tyramide SuperBoost™ Kit (goat anti-mouse IgG, Invitrogen, B40912) was used to amplify the signal. All slides were mounted with ProLong Gold Antifade reagent with DAPI (Invitrogen, P36931). Images were obtained using Leica Software on a Leica SPE confocal microscope and analyzed in ImageJ.

### Hematoxylin & Eosin staining and image analysis

Paraffin-embedded tissue sections were incubated in xylene (twice, 10 minutes each), then rehydrated through graded ethanol (3 minutes each), and finally washed with water twice for 5 minutes. Next, tissue sections were stained with hematoxylin for 30 seconds, washed immediately with running water for 5 minutes, and then stained with eosin for 1 minute. After a short dip in water, tissue sections were dehydrated with graded ethanol (70% for 2 min, 95% for 2 min, 100% for 2 min twice, and xylene for 5 min twice). Tissue sections were mounted in Organo/Limonene mounting medium and dried overnight. H & E-stained tissue sections were scanned using a Fast Scanner at the Imaging Facility at Albert Einstein College of Medicine. Scanned images were analyzed for metastasis by CaseViewer v2.4.

### Western blot analysis

Total proteins were extracted using RIPA buffer (50 mM Tris pH 7.4, 1 mM EDTA, 150 mM NaCl, 1% Triton X-100, 0.25% sodium deoxycholate, o.1% SDS, 1% NP40). Samples were boiled in Laemmli sample buffer for 15 minutes at 95°C. 10-12% SDS–PAGE gels were run in running buffer (25 mM Tris, 190 mM glycine, 0.1% SDS) and transferred to PVDF membranes in transfer buffer (25 mM Tris, 190 mM glycine, 20% methanol). Membranes were blocked in 5% milk or BSA in TBST (Tris-buffered saline with 0.1% Tween-20) buffer. Membranes were incubated with primary antibodies overnight at 4°C. Membranes were washed in TBST buffer and incubated with HRP-conjugated secondary antibodies at room temperature for 1 hour. Western blots were developed with Pierce ECL Western Blot Detection substrate, and images were captured with the Invitrogen iBright 1500 imaging system.

### RT-qPCR

Total RNA was extracted using the RNeasy Plus Micro kit (Qiagen, 74034). Reverse transcription was performed using the iScript gDNA clear cDNA synthesis kit (BioRad, 1725035). Relative expression levels of RNA were determined using PowerUP SYBR^TM^ Green master Mix (Applied Biosystems, A25776) by a QuantStudio 3 Real-Time PCR System (Applied Biosystems, A28567). Gene expression levels were normalized to the *B2m* housekeeping gene. Primers used for RT-qPCR are listed in the **supplementary information-Table 2**.

### RNA-sequencing and data analysis

Total RNA was extracted using the Qiagen RNeasy Plus Micro kit. RNA samples QC, library preparation and RNA sequencing were performed at GENEWIZ, LLC. (South Plainfield, NJ, USA). Concentration of RNA samples was estimated by Qubit 2.0 Fluorometer (Life Technologies) and RNA integrity (RIN) was checked by Agilent TapeStation 4200 (Agilent Technologies). RNA samples were enriched for Poly-A mRNA using NEBNext Poly(A) mRNA Magnetic Isolation Module (New England Biolabs, Ipswich, MA, USA), and RNA-seq library was prepared using the NEBNext Ultra II RNA Library Prep Kit for Illumina following the manufacturer’s recommendations. The samples were sequenced in a 2 x 150 bp paired-end configuration using the Illumina NovaSeq platform. After quality controls, the paired-end reads were aligned to the mouse reference genome (mm39) using STAR (v2.7.9a).^49^ Read counts and transcripts per million (TPM) were quantified with RSEM (v1.3.3)^50^ based on gene annotations from the Ensembl database (v105). Genes with TPM ≥ 1 in at least one sample of a comparison were retained for the downstream analysis. Differential expression analysis was performed using DESeq2 (v1.38.3).^51^ A false discovery rate (FDR) < 0.05 was used to identify differentially expressed genes (DEGs).

### Human Data Analysis

We derived Zfp281-associated transcriptional signatures from differential expression analyses of mouse perturbation experiments that compared knockout (KO) or overexpression (OE) with wild-type samples using DESeq2 (adjusted P<0.05). To enable analysis in human datasets, mouse genes were mapped to human orthologs using Ensembl BioMart, and genes without a mapped human ortholog were excluded. Upregulated and downregulated genes from each perturbation defined four signatures: KO up, KO down, OE up, and OE down. Two human expression datasets were analyzed. The METABRIC dataset (1980 samples) contains primary tumors annotated by the IC10 subtypes. The DCIS dataset (353 samples) includes primary and pre-invasive samples with known recurrence status.

Signature genes were intersected with the genes present in each dataset. Single-sample gene set variation analysis (GSVA) was then performed using GSEApy on log2-transformed TPM values, with a minimum gene set size of 5 genes, to obtain enrichment scores for each signature in each sample. A composite Zfp281 activity score was defined as follows: (𝐾𝑂 ↓ − 𝐾𝑂 ↑) + (𝑂𝐸 ↑ − 𝑂𝐸 ↓) using GSVA enrichment scores. When a signature pair was incomplete, the missing component was excluded from the calculation. For example, if KO up genes were absent, the score was computed using only the OE component. Group differences were evaluated using two-sided Mann-Whitney U tests. Where appropriate, P-values were adjusted using the Benjamini–Hochberg false discovery rate procedure.

### Tumor doubling time

Doubling time was calculated using the following formula: DT (days) = ln(2) x Δt / ln(M2/M1). Δt is the time (days) between tumor measurements; M1 and M2 are the initial tumor size and the last tumor size, respectively.

### Statistical analysis

Unless otherwise specified, three independent *in vitro* experiments were performed, and *in vivo* tissue analyses were conducted with at least five mice per condition. Statistical analyses were done using Prism software, and differences were considered significant if P < 0.05. The absence of P-values indicates insignificant differences.

## DATA AVAILABILITY

Bulk RNA-sequencing data have been deposited with the GEO under the accession number GSE325780. All other raw data generated in this study are available upon request from the corresponding authors. We additionally utilized publicly available METABRIC data (Project SynID: syn1688369).

## AUTHORS’ CONTRIBUTIONS

**D.K. Singh:** conceptualization, validation, investigation, visualization, methodology, formal analysis, writing-original draft, writing-review, and editing. **H. Zhou:** methodology, validation, investigation. **N. Sherpa:** methodology, investigation. **X.Y. Zheng:** formal analysis. **A. Lomakin:** formal analysis. **P. Razghandi:** methodology. **X. Huang:** methodology. **R. Kadamb:** methodology. **S. Shukla:** formal analysis. **L.E. Valencia Salazar:** methodology. **D. Entenberg:** supervision. **D. Zheng:** data curation, supervision. **C. Curtis:** resources, data curation, supervision. **J. Wang:** conceptualization, resources, funding acquisition, supervision, writing-review, and editing. **J.A. Aguirre-Ghiso:** conceptualization, resources, funding acquisition, supervision, writing-review, and editing.

## ACKNOWLEDGEMENTS

This work has been supported by the Department of Defense (DoD, BC200469 to J.W. and BC200469P1 to J.A.A-G). J.W. is supported by grants from the National Institute of Health (NIH) /National Cancer Institute (NCI) (HD114122, HD117611, and CA285299). J.A.A-G. is supported by grants from the NIH /NCI (CA109182, CA284085, CA301643, CA253977, and CCSG P30 CA013330-52), and The Gurwin Foundation and the Garfunkel, Spiegel, and Spatz families for their generous support. J.A.A-G. is also supported by a DOD CDMRP FY23 BCRP FL2 PSPS (BC230537-HT94252410078) and an Aging and Cancer grant from the Samuel Waxman Cancer Research Foundation and The Mark Foundation for Cancer Research. L.E.V.S was funded by a Medical Scientist Training Program grant from the National Institute of General Medical Sciences (NIGMS) of the NIH under award number T32GM149364 to the Albert Einstein College of Medicine MD-PhD Program. H&E and live cell imaging were performed at the Advanced Imaging Facility at the Albert Einstein College of Medicine, which is supported by a Cancer Centre grant (P30CA013330) and a Shared Instrumentation Grant (S10OD026852-01A1), with assistance from Vera Desmarais.

---

**Supplementary Figure 1.**
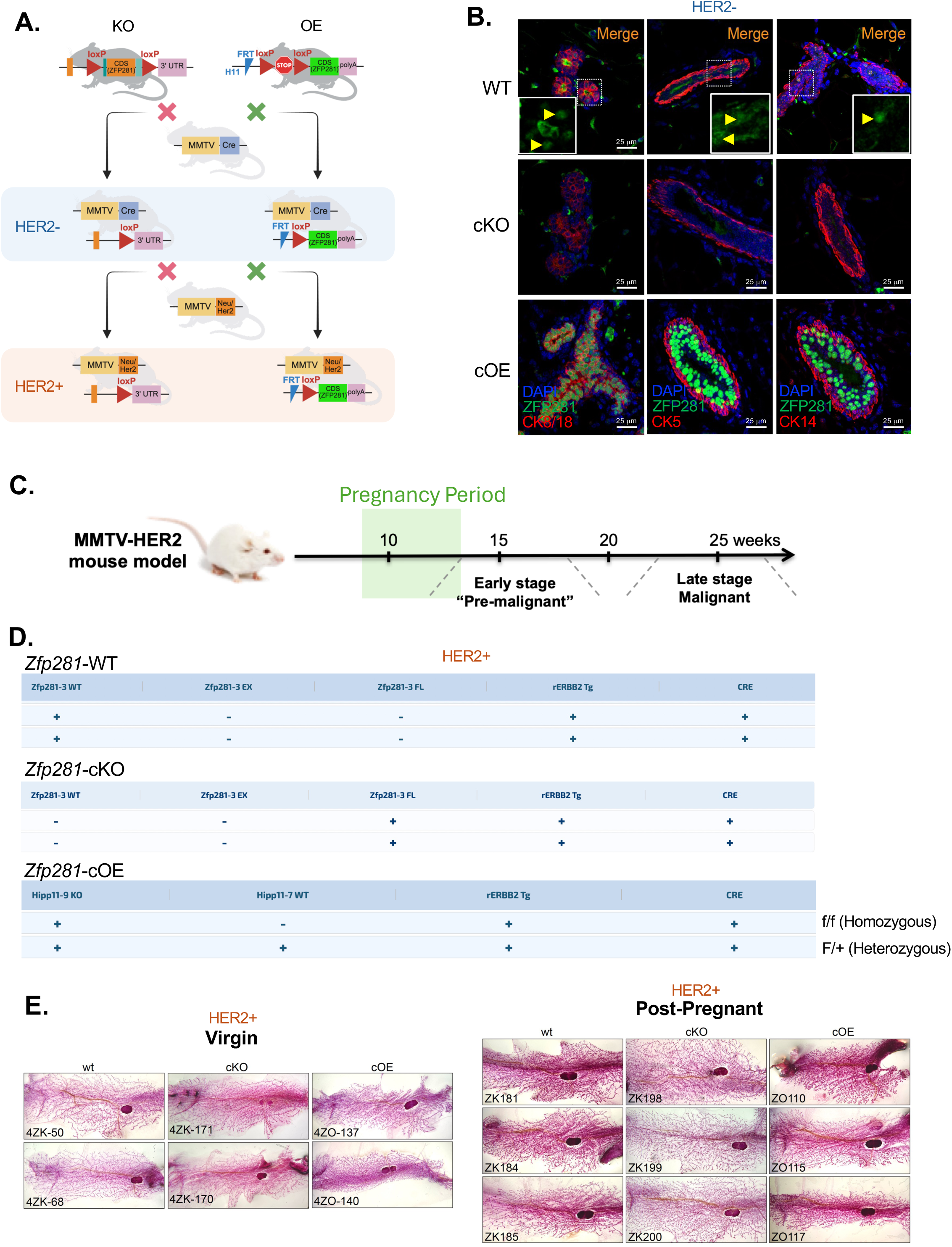
**A.** A schematic showing the development of conditional knockout and conditional overexpression of Zfp281 in the mammary gland of the mouse. **B.** Confirmation of ZFP281 KO and OE in the normal (HER2-) mammary gland of ZFP281 WT, cKO and cOE mice by immunofluorescence staining. Luminal cells of mammary ducts expressing ZFP281 in the WT mice are marked with yellow arrows in the enlarged images. **C.** A schematic showing the timeline for the pregnancy, early lesion stage, and primary tumor development in the MMTV-HER2 breast cancer mouse model. **D.** Genotyping data (performed at TransnetYX) confirming the KO and OP of *Zfp281* in the MMTV-HER2 mouse model. EX: Exon, FL: Floxed, Tg: Transgene. Zfp281-3FL positive, rERBB2 Tg positive and Cre positive are Zfp281 KO mice. Hipp11-9 KO positive, Hipp11-7Wt negative, rERBB2 Tg positive, and Cre positive are Zfp281 OE (homozygous) mice, while those positive for Hipp11-7WT are also Zfp281 OE (heterozygous). **E.** Mammary glands whole mount in MMTV-HER2 ZFP281 WT, cKO, and cOE virgin and post-pregnant mice.

**Supplementary Figure 2.**
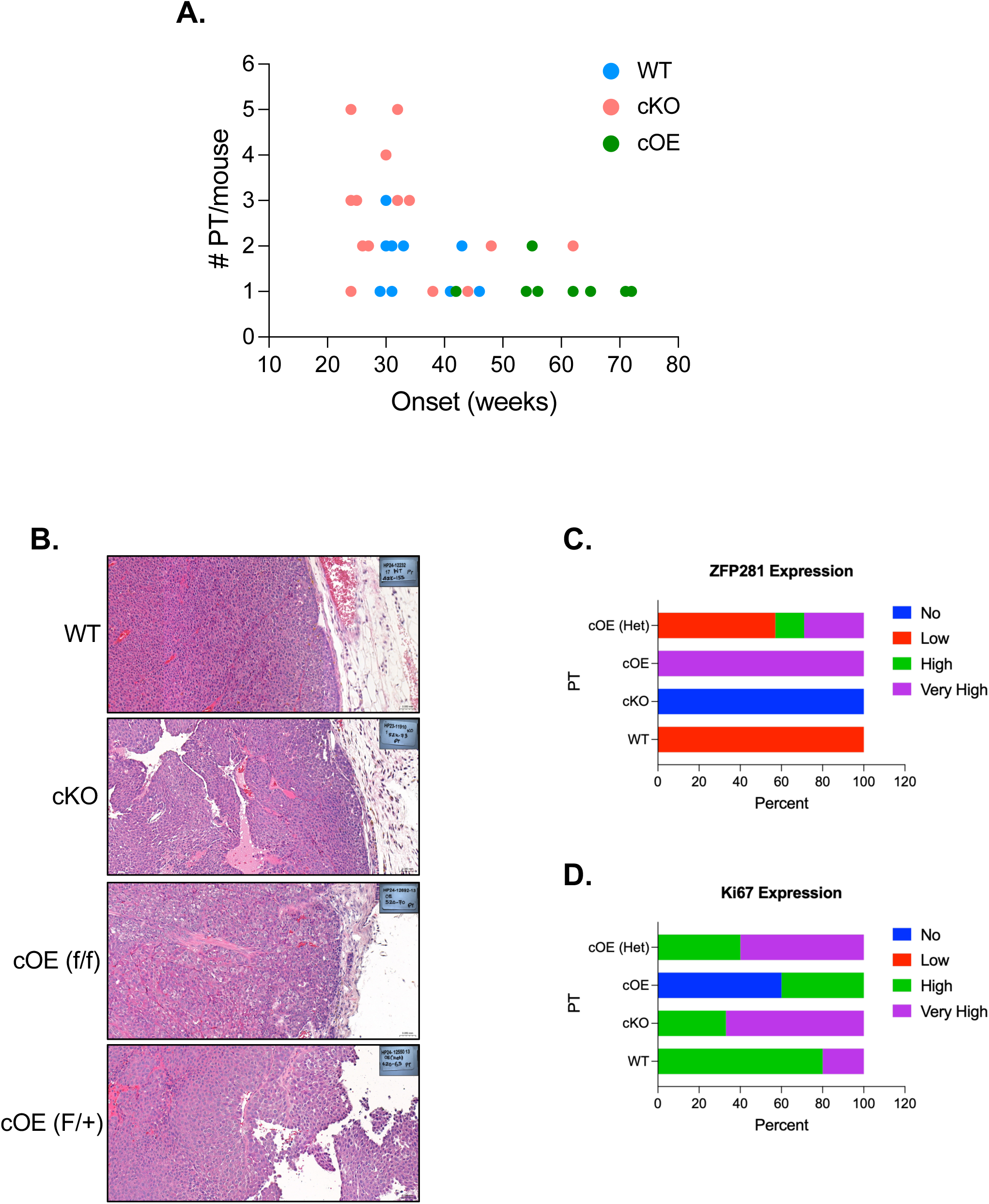
**A.** The graph shows the correlation between the number of primary tumors and the timing of onset (detection of the first PT). **B.** H & E staining of primary tumors. **C-D.** The graph shows the stratification of ZFP281 and Ki67 expression in the PT developed in the MMTV-HER2 ZFP281 WT, cKO and cOE (heterozygous and homozygous).

**Supplementary Figure 3.**
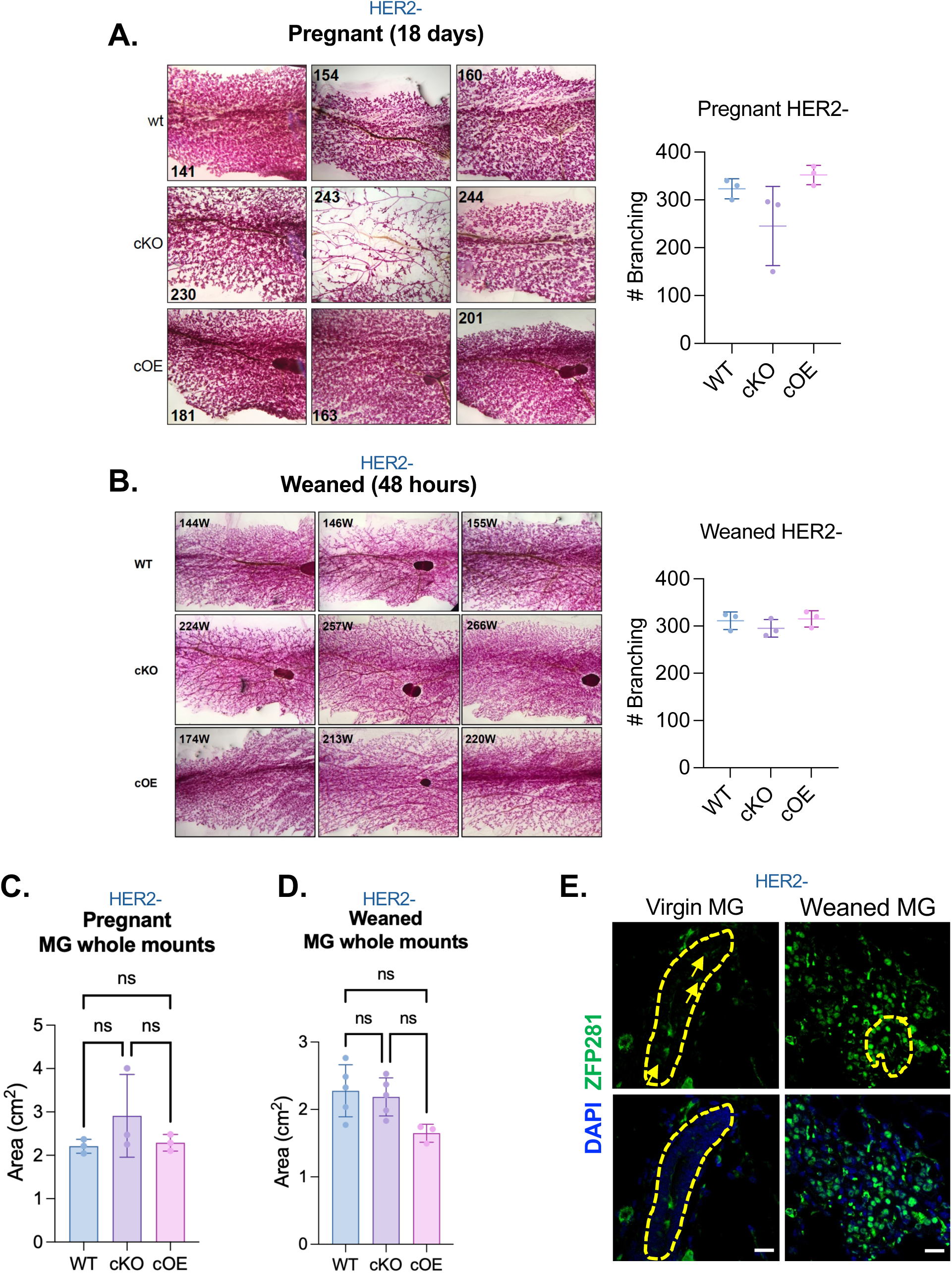
**A-B.** Whole-mount mammary glands and their quantification in MMTV-Cre (HER2-) ZFP281 WT, cKO, and cOE pregnant and post-weaned (48 hours after weaning) mice. **C-D.** Mammary gland size quantifications in pregnant and post-weaned mice. **E.** Expression of ZFP281 in the mammary glands of ZFP281 WT virgin and post-weaned mice detected by immunofluorescence staining. Yellow boundaries highlight the mammary ducts.

**Supplementary Figure 4.**
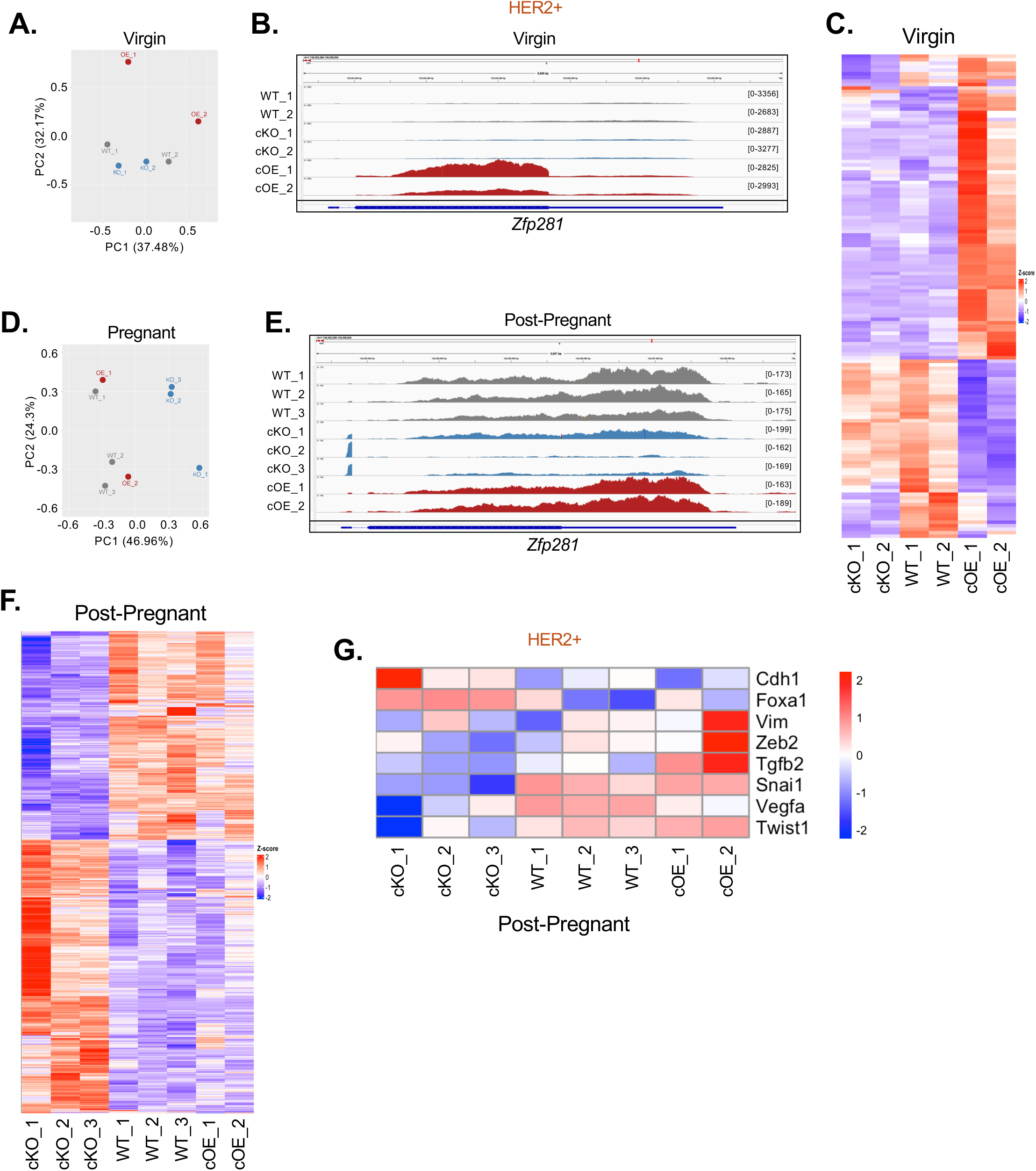
**A.** PCA plot of bulk RNA-sequencing in MMTV-HER2 ZFP281 WT, cKO and cOE virgin mice. **B.** Snapshot of Zfp281 expression from RNA-sequencing in virgin mice. **C.** Heatmap showing comparison of differentially expressed genes between ZFP281 WT, cKO, and cOE mice at the virgin stage. **D.** PCA plot of bulk RNA-sequencing in MMTV-HER2 ZFP281 WT, cKO, and cOE post-pregnant mice. **E.** Snapshot of Zfp281 expression from RNA-sequencing in post-pregnant mice. **F.** Heatmap showing comparison of differentially expressed genes between ZFP281 WT, cKO, and cOE mice at the post-pregnant stage. **G.** Expression of EMT genes (by RNA-seq) in the post-pregnant mice.

**Supplementary Figure 5.**
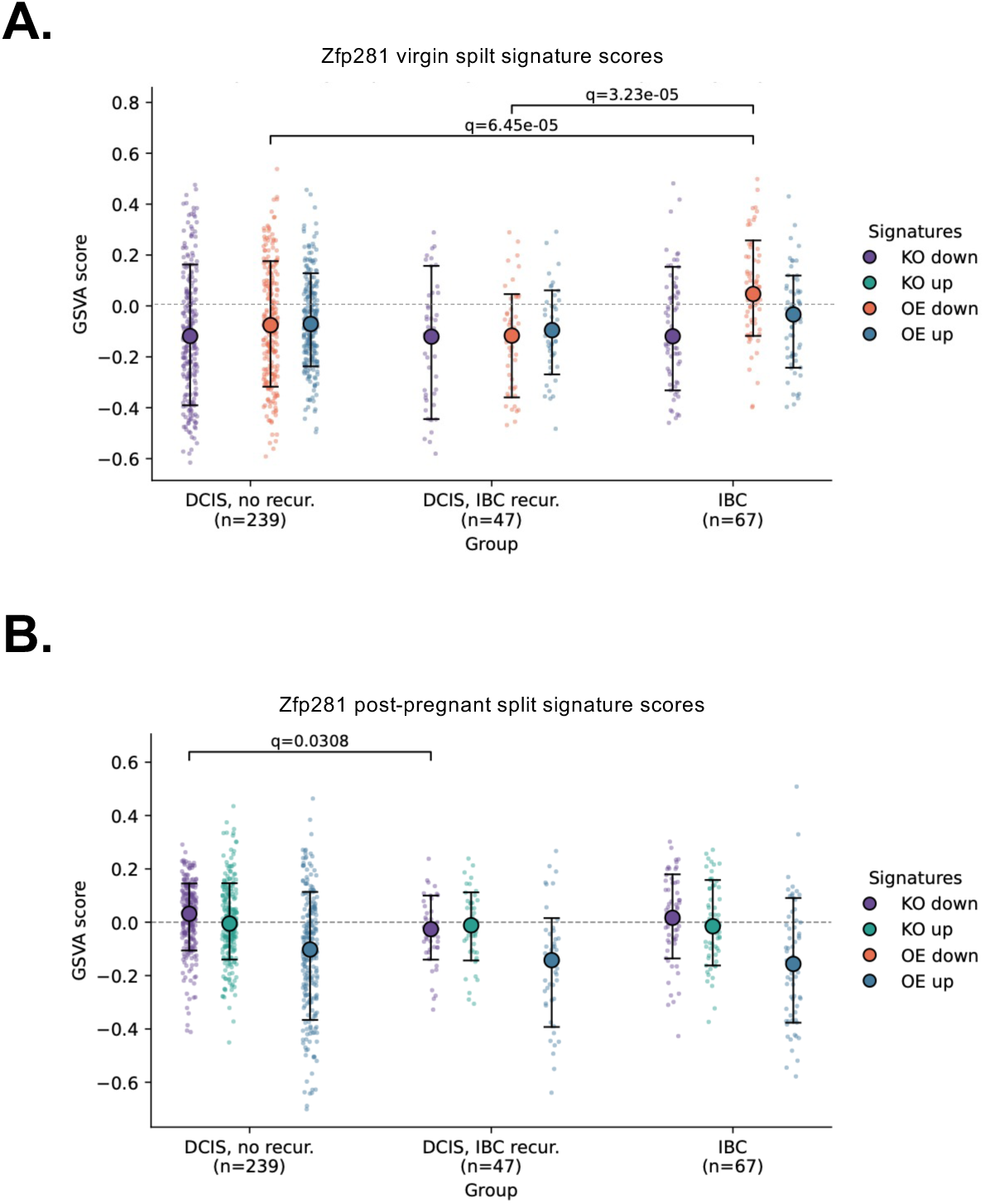
**A-B.** Genes differentially expressed upon ZFP281 KO and OE were used as signatures for GSVA in the DCIS and IBC cohorts. KO-up in virgin and OE-down in post-pregnancy were not used as signatures, as they contain only 1 and 8 genes, respectively.

