## Supplementary Information for "Enforced ZFP281 expression delays breast cancer initiation and can provide lifelong protection against breast cancer metastasis"

### Mouse model generation and genotyping methods

1. Generation of *Zfp281* conditional knockout (cKO) mouse strain

We generated a *Zfp281* conditional knockout (cKO) mouse strain by using CRISPR/Cas9 and Cre/loxP systems.[1] To construct the *Zfp281* conditional allele, we used the "two-donor floxing method", which uses two sgRNAs and two single-stranded oligonucleotides (ssODNs) containing 34 bp loxP sites to flank a targeted critical exon.[2,3] *The zfp281* gene has a unique 2-exon structure, with Exon2 encoding the full-length protein and Intron1 containing a highly GC-rich sequence, posing a potential challenge for loxP insertion. Accordingly, we knocked in one loxP site (loxP1) at Intron1 and a second loxP site (loxP2) at the position immediately downstream of the stop codon to minimize the possible disruption of 3' UTR. We designed sgRNAs and donor ssODNs with the loxP sequence flanked by 60 bp of homology arms. To facilitate the detection of correct targeting, each ssODN was engineered to contain a BamHI restriction site. The Cas9 mRNA, sgRNAs, and ssODNs were injected into 150 zygotes, and the *in vitro* developed blastocysts were then transplanted into the uterus of pseudo-pregnant females to generate CRISPR founder mice. We obtained 23 (out of 150) founder pups in total and then genotyped mouse tails using next-generation sequencing (NGS). Four out of 23 mice contained both loxP insertions at the same allele, and the knock-in sequence was confirmed with correctly targeted insertions. We backcrossed the four founder mice with wild-type (WT) C57BL/6 mice to transmit and segregate the KI allele among the F1 *Zfp281*$^{F/+}$ mice, then intercrossed the *Zfp281*$^{F/+}$ male and female mice to obtain the homozygous *Zfp281*$^{F/F}$ mice. The correct integration of loxP sites was further confirmed by Sanger sequencing. The adult *Zfp281*$^{F/F}$ mice were normal in body weight and fertility compared to the WT C57BL/6 mice. Thus, we successfully generated the first *Zfp281* conditional knockout (cKO) mouse model by overcoming the potential loxP insertion hurdle associated with the unique 2-exon structure and the GC-rich Intron 1 of the *Zfp281* gene. PCR genotyping primers: 5'-CGGAACCCgtaagtgtgg-3'; 5'-GAACTTTGCACAAGCATTCG-3'; 5'-AGTTGCATTGAAAGGGCATT-3'; WT(+): with a band of 322 bp; Flox (F): one band with 362 bp; KO (-): one band with 270 bp.

2. Generation of *Zfp281* conditional overexpression (cOE) mouse strain

*Zfp281* conditional overexpression (cOE) mouse was generated by CRISPR knockin the Lox-STOP-Lox-Zfp281 (LSL) cDNApA cassette into the Hipp11 (H11) locus (note: H11 replaces Rosa26). The gRNA (GAACACTAGTGCACTTATCCTGG) to mouse Hipp11 locus, the donor vector containing "Rox-CAG promoter-loxP-Kozak-EGFP-3*polyA-loxP-Kozak-mouse Zfp281 CDS-polyA-Rox" cassette, and Cas9 mRNA were co-injected into fertilized mouse eggs to generate targeted conditional knock-in offspring. F0 founder animals were identified by PCR followed by sequence analysis and were bred to wild-type mice to test germline transmission and F1 animal generation. Then, Inter-cross heterozygous targeted mice to generate homozygous targeted mice (L/L). PCR primers: 5'-CTCTACTGGAGGAGGACAAACTG-3'; 5'-GTCTTCCACCTTTCTTCAGTTAGC; 5'-GCATCTGACTTCTGGCTAATAAAG-3'; Homozygous (L/L): one band with 785 bp; Heterozygous (L/+): two bands with 785 bp/ 519 bp; WT: one band with 519 bp. Upon Cre excision, L/+ will have 2 (endogenous) + 1 (H11) = 3 alleles of Zfp281. Thus, cOE L/L will end up with 4 x Zfp281 alleles upon Cre excision.

MMTV-Cre (Line D) was obtained from JAX with Stock No. 003553: both Zfp281 cKO (F/F) and cOE homozygous targeted mice (L/L) were bred with an MMTV-Cre mouse to get final F/F-MMTV-Cre or L/L-MMTV-cre mice. We carried out the genotyping followed JAX protocol(s). For Cre primers: Transgene Forward (Cre-F): GCG GTC TGG CAG TAA AAA CTA TC; Transgene Reverse (Cre-R): GTG AAA CAG CAT TGC TGT CAC TT; Internal Positive Control Forward: CTA GGC CAC AGA ATT GAA AGA TCT; Internal Positive Control Reverse: GTA GGT GGA AAT TCT AGC ATC ATC C; Transgene=~100 bp; Internal positive control=324 bp.

**Table 1. Antibodies**

| Name | Source | Catalogue # |
|---|---|---|
| ZFP281 | Santacruz Biotechnology | sc-166933 |
| CK8/18 | Bioss antibody | BSM-52419R |
| CK5 | Abcam | ab52635 |
| CK14 | Abcam | ab7800 |
| HER2 | Sigma | HPA001383 |
| Ki67 | Invitrogen | 14-5698-82 |
| $\alpha$-Tubulin | Santacruz Biotechnology | sc-32293 |

**Table 2. Primers**

| Primer | Forward (5'-3') | Reverse (5'-3') |
|---|---|---|
| *Zfp281* | GCAGTGCGTGTTATCCTCCT | ATGCTCACCAAGGACTGCAA |
| *E-Cadherin* | AACCCAAGCACGTATCAGGG | GAGTGTTGGGGGCATCATCA |
| *N-Cadherin* | CACTGCCATTGATGCGGATG | TGCCACAGTGATGATGTCCC |
| *Vimentin* | TTCTCTGGCACGTCTTGACC | GCTTGGAAACGTCCACATCG |
| *Twist1* | CCCACACCTCTGCATTCTGAT | CAGTGGCTGATTGGCAAGAC |
| *Snail* | CTCCAAACCCACTCGGATGT | AGCCAGACTCTTGGTGCTTG |
| *Zeb1* | ACACAAGCGAGAGGATCATGG | ATAATTTGTAACGTTATTGCGCCG |
| *Myc* | GGTGTCTGTGGAGAAGAGGC | TTGTGCTGGTGAGTGGAGAC |
| *Gata3* | CCATTACCACCTATCCGCCC | TTCACACACTCCCTGCCTTC |
| *Klf4* | CCGACTAACCGTTGGCGT | CGGGTTGTTACTGCTGCAAG |
| *β-Actin* | CCAGCCTTCCTTCTTGGGTAT | GGGTGTAAAACGCAGCTCAG |
| *B2m* | CTCGGTGACCCTGGTCTTTC | GGATTTCAATGTGAGGCGGG |
